# Molecular switch mediates the glucocorticoid receptor transition from tumor suppressor to oncogene in the prostate

**DOI:** 10.1101/2025.08.20.670078

**Authors:** Johannes Hiltunen, Niina Aaltonen, Heini Sohlberg, Laura Kemppi, Konsta Kukkonen, Tapio Visakorpi, Ville Paakinaho

**Affiliations:** Institute of Biomedicine, University of Eastern Finland, Kuopio, Finland; Faculty of Medicine and Health Technology, Tampere University and Tays Cancer Centre, Tampere University Hospital, Tampere, Finland; Fimlab Laboratories Ltd, Tampere University Hospital, Tampere, Finland

**Keywords:** chromatin, GATA2, glucocorticoid receptor, p63, prostate cancer

## Abstract

Glucocorticoids are widely used to alleviate inflammation and treatment-related side effects in prostate cancer (PCa), particularly to counteract abiraterone-induced cortisol suppression. However, the glucocorticoid receptor (GR) exhibits a dual role exerting tumor-suppressive effects by inhibiting early-stage PCa cell proliferation, while also promoting oncogenic progression by mediating antiandrogen resistance. The molecular mechanisms underlying this functional dichotomy have remained elusive and poorly characterized. Using genome-wide analyses and CRISPR-based genome editing, we identified the tumor protein p63 as a key mediator of GR’s tumor-suppressive chromatin activity and maintenance of basal epithelial cell identity. Loss of p63 reprograms GR activity toward an oncogenic state, marked by basal-to-luminal transition, enhanced cell migration, invasion, and altered morphology accompanied by increased epithelial-mesenchymal transition markers. This functional shift is driven by elevated expression of transcription factors GATA2 and FRA1, which remodel GR’s chromatin binding and transcriptional output to activate oncogenic signaling pathways. Together, our findings uncover a molecular switch that governs the dual role of GR in PCa, establishing transcription factor crosstalk as a critical regulator of GR-driven oncogenic reprogramming and cellular plasticity. These findings establish a framework for understanding glucocorticoid-induced tumor suppression and highlight GATA2 and FRA1 as potential targets to mitigate GR-mediated resistance in PCa.

## Introduction

Prostate cancer (PCa), one of the most prevalent cancers worldwide ^1^, is primarily driven by signaling through the androgen receptor (AR) ^2,3^. AR belongs to the steroid receptor subfamily of nuclear receptors, which also includes estrogen receptor (ER) and glucocorticoid receptor (GR). These receptors are ligand-dependent transcription factors (TFs) that, upon hormone binding, translocate to the nucleus, bind chromatin, and regulate gene expression. This regulation is mediated through coordinated recruitment of TFs and coregulators with histone-modifying and chromatin-remodeling functions ^4,5^.

Because AR-driven gene regulatory programs are central to PCa development and progression, inhibition of AR signaling remains the cornerstone of PCa therapy. Depending on disease stage, treatment strategies include androgen deprivation therapy (ADT), antiandrogens such as enzalutamide and darolutamide, and androgen biosynthesis inhibitors like abiraterone ^2,3^. Although these therapies are initially effective, many tumors eventually develop resistance to AR-targeted treatments. In this resistant state, cancer progression is often sustained by alternative TFs ^6^. Notably, TFs such as FOXA1, GATA2, and SOX2 can reprogram chromatin accessibility and promote lineage plasticity, enabling continued tumor growth in the absence of AR activity ^7–9^. Moreover, these TFs can drive antiandrogen resistance by modulating the activity of other steroid receptors, including GR ^10,11^.

Glucocorticoids—the activating ligands of GR—have traditionally been used as adjuvant therapy in PCa to alleviate treatment-related side effects ^12^. However, GR is now recognized as a context-dependent oncogene in PCa, capable of mediating drug resistance by compensating for the loss of AR signaling under antiandrogen therapy ^13^. While the pro-tumorigenic role of GR in therapy-resistant PCa has been well documented by us and others ^10,14,15^, its function during earlier stages of the disease remains elusive. Notably, early studies suggested a tumor suppressive role for GR in primary PCa, where it inhibited proliferation and oncogenic signaling pathways ^16^. These findings point to a striking functional shift in GR activity—from tumor suppressor to oncogene—throughout disease progression. However, the molecular mechanisms underlying this transition remain poorly understood. Given GR’s ubiquitous expression, this switch is likely governed by changes in the transcriptional landscape and signaling networks driven by other TFs.

It is well established that the tumor suppressive or oncogenic activity of a TF can be modulated by TF crosstalk, as well as by lineage and epigenetic plasticity ^7,17,18^. For steroid receptors, the clearest examples of this come from breast cancer. Both GR and AR exhibit tumor suppressive roles in ER-positive breast cancers, while exerting oncogenic functions in ER-negative subtypes ^4,19,20^. This highlights how the presence or absence of ER signaling can dramatically reshape the functional consequences of other steroid receptors. Moreover, this context-dependent shift is not limited to interactions among steroid receptors. For example, AR’s tumor-suppressive role in ER-positive breast cancer can be reprogrammed into an oncogenic function through activation of the JAK/STAT pathway ^21^. These findings underscore the capacity of distinct TF networks and signaling pathways to serve as molecular switches, governing the tumor suppressor-to-oncogene transition of steroid receptors.

To investigate the transition of GR from a tumor suppressor to an oncogene in the prostate, we examined its transcriptional activity across various PCa cell lines and immortalized normal prostate epithelial cells. We found that both the GR-regulated transcriptome and its cistrome differ markedly between normal and cancerous prostate cells, regardless of PCa cells’ AR status. In normal prostate cells, GR binding sites are uniquely co-occupied by the basal epithelial TF p63, a member of the p53 family. Loss of p63 diminishes GR’s anti-proliferative effects and instead promotes GR-driven cell invasion. This phenotypic switch is associated with increased GATA2 and FRA1 activity. In the absence of p63, GR’s transcriptional output is reprogrammed from tumor suppressive to oncogenic pathways. Together, these findings suggest that GR’s oncogenic transition in prostate cancer is governed by dynamic changes in transcription factor crosstalk.

## Materials and Methods

### Cell culture and treatments

RWPE1 derived cells (ATCC, # CRL-3607, RRID:CVCL_3791) and RWPE1-ARc15 (RWPE1-AR) cells ^22^ were cultured in Keratinocyte SFM (Thermo-Fischer, #17005042) supplemented with BPE (50 µg/ml) and EGF (5 ng/ml) and 1 U/µl penicillin and 1 µg/ml streptomycin (Gibco, #15140122). DU145 cells (DSMZ, #ACC261, RRID:CVCL_0105) were cultured in MEM (Thermo-Fisher, #10370047) supplemented with 10% FBS (Gibco, #11573397), 2 mM l-glutamine (Gibco, #A2916801), and 1 U/µl penicillin and 1 µg/ml streptomycin (Gibco, #15140122). PC3 cells (ATCC, # CRL-1435, RRID:CVCL_0035) were cultured in Ham’s F12-K medium (Gibco, #21127022) supplemented with 10% FBS (Gibco, #11573397), 1 U/μl penicillin and 1 μg/ml streptomycin (Gibco, #15140122). PNT1A cells (Sigma, #95012614, RRID:CVCL_2163) were cultured in RPMI-1640 medium (Gibco, #1187509) supplemented with 10% FBS (Gibco, #11573397), 2 mM l-glutamine (Gibco, #A2916801), and 1 U/µl penicillin and 1 µg/ml streptomycin (Gibco, #15140122). For experiments DU145, PC3, and PNT1A were cultured in experiment medium containing 5% charcoal stripped FBS (Gibco, #10270106), for minimum of 2 days before the start of hormone treatment. Dexamethasone (Dex) (Sigma-Aldrich, #D4902) and dihydrotestosterone (DHT) (Steraloids, #A2570-000) treatments were conducted with 100 nM concentration. The cells were regularly tested for mycoplasma contamination.

### Generation of knockout cell lines using CRISPR-Cas9

Predesigned Alt-R CRISPR-Cas9 crRNA (IDT) (Supplementary Table S1) recognized the DNA binding domain in exon 5 of *TP63* gene. Alt-R CRISPR-Cas9 negative control crRNA (IDT, # 1072544) was used as control. crRNA and tracrRNA (IDT, #1072533) were combined to create each guide RNA (gRNA). *TP63*-specific gRNA was predicted to induce breaks before the key amino acids critical for Zinc Finger, DNA binding, and Intradimer functionality ^23^. Alt-R *S.p.* Cas9 Nuclease V3 (IDT, #1081058), was diluted to 1 µM concentration in Cas9 working buffer and was incubated with gRNA oligos to form ribonucleoprotein (RNP) complex. Generation of gRNAs, RNPs and cationic lipid delivery were performed according to manufacturer’s instructions (IDT). RNP complexes were reverse-transfected into RWPE1 cells using Opti-MEM (Thermo Fisher, #31985070) and Lipofectamine RNAiMAX Transfection Reagent (Invitrogen, #13778150). Initial gRNAs functionality was tested with Alt-R Genome Editing Detection Kit (IDT, #1075932) and Agilent Bioanalyzer 2100 DNA 1000 Analysis kit (Agilent, #5067–1504). After 48 h exposure of RWPE1 cells to RNPs, cells were collected and plated in 96-well plate with serial dilutions to obtain monoclonal cultures, with monoclonality verified by microscopy. After the expansion of monoclonal cultures, the absence of p63 (p63KO) was verified with immunoblotting. For verification of genome editing, genomic DNA was isolated using DNA Extraction Solution (Epicentre, #QE09050), and Sanger sequenced (Macrogen Europe) (Supplementary Table S1).

### Antibodies

Primary antibodies: anti-GR (Cell Signaling Technology, #12041, RRID:AB_2631286), anti-α tubulin (Santa Cruz Biotechnology, #sc-5286, RRID:AB_628411), anti-p53 (Abcam, #ab1101, RRID: AB_297667), anti-p63α (Cell Signaling Technology, #13109, RRID:AB_2637091), anti-FRA1 (Cell Signaling Technology, #5281, RRID:AB_10557418), anti-GATA2 (Cell Signaling Technology, #79802, RRID:AB_3105938), anti-N-Cadherin (Cell Signaling Technology, #13116 , RRID:AB_2687616), anti-TWIST1 (Cell Signaling Technology, #69366, RRID:AB_2891135), anti-Vimentin (Cell signaling technology, #5741, RRID:AB_10695459), anti-GAPDH (Thermo Fisher, PA1-987, RRID:AB_2107311), anti-H3K27ac (Active Motif, #39133, RRID:AB_2561016), anti-AR ^24^. Secondary antibodies: goat anti-rabbit (Invitrogen, #G-21234, RRID:AB_1500696), goat anti-mouse (Invitrogen, #62-6520, RRID:AB_2533947).

### Immunoblotting

Protein isolation and separation was done as before ^25^. Primary antibodies: anti-GR (1:1000), anti-p63α (1:1000), anti-GATA2 (1:1000), anti-N-Cadherin (1:1000), anti-TWIST1 (1:1000), anti-Vimentin (1:1000), anti-AR (1:10 000). Anti-tubulin antibody (1:3000) or anti-GAPDH antibody (1:5000) was used as a loading control. Secondary antibodies (1:10000): anti-mouse (tubulin), anti-rabbit (GR, p63, GATA2, N-Cadherin, TWIST1, Vimentin, GAPDH). The detection was done using Pierce ECL Western Blotting Substrate kit (Thermo Fisher, #32106) with Bio-Rad ChemiDoc Imager.

### RNA extraction and RNA-seq

For RNA extraction, the cells were plated on 6-well plates (PC3, 200k cells per well; DU145, 300k cells per well; RWPE1, 250k cells per well). The cells were treated with vehicle (0.1% EtOH), or 100 nM Dex for 6 h before the start of RNA extraction. For RNA interferences, the cells were reverse siRNA transfected using Opti-MEM (Thermo-Fisher, #31985070) and Lipofectamine RNAiMAX Transfection Reagent (Invitrogen, #13778150) according to manufacturer’s instructions. SiRNAs were used at 60 nM concentration: ON-TARGET plus Non-targeting Control SMARTpool (siNON, #D-001810–10), ON-TARGET plus Human GATA2 SMARTPOOL (siGATA2, L-009024-02-0020). Immunoblotting was used to verify siRNA effectiveness as above. RNA was extracted using TriPure Isolation Reagent (Roche, #1166715700), and Monarch Total RNA Miniprep Kit (New England BioLabs, #T2010S). RNA sequencing (RNA-seq) libraries were generated with NEBNext Poly(A) mRNA Magnetic Isolation Module (New England BioLabs, #E7490) and NEBNext Ultra II Directional RNA Library Prep Kit (New England BioLabs, #E7765) according to manufacturer’s instructions using 2 µg of purified RNA. Library quality was measured with Agilent 2100 Bioanalyzer using DNA 1000 analysis kit (Agilent, #5067-1504). Two to three replicates were then sequenced with Illumina NextSeq 500 (75SE) or Illumina NextSeq 2000 (100SE).

Human hg38 genome alignment was done using STAR 2.7.9a (RRID:SCR_004463) ^26^ with default settings allowing max 10 mismatches and max 10 multi-mapped reads. Differential expression was assessed with DESeq2 (RRID:SCR_015687) ^27^ and HOMER (4.1.1) (RRID:SCR_010881) ^28^. For visualization genes were normalized transcript per million (TPM). Differentially expressed genes were defined with log_2_ fold change ± 0.5 and FDR < 0.05 between specific conditions. DESeq2 data and TPM values are presented in Supplementary Table S2. Z-score pathway average was calculated by taking an average from single pathway gene Z-scores. Metascape pathway enrichment analysis (RRID:SCR_016620) ^29^ was used for comparison of specific gene clusters. Hallmark, CORUM, and WIKI pathway gene sets were used with criteria *p* < 0.01, at least three gene overlap per gene set, and > 1.5 pathway enrichment. For genes downregulated by siGATA2 (siGATA2 DN), log_2_ fold change >1 and FDR < 0.05 requirement were utilized. For Dex-regulated siGATA2 DN genes, siGATA2 led to loss of Dex-regulation wherein the target genes no longer met the above set criteria.

### ChIP-seq and ATAC-seq

For chromatin immunoprecipitation (ChIP), the cells were plated on 10-cm plates (PC3, 1M cells per plate; DU145, 2M cells per plate; RWPE1, 2M cells per plate; PNT1A, 1.5M cells per plate). K-7174 treatments were performed 24 h before sample crosslinking. SiRNA treatments with ON-TARGETplus 2.0 Human TP63 siRNA (siTP63, L2-003330-01-0010) and ON-TARGETplus 2.0 Non-targeting Control Pool (siNON, D2-001810-10) were performed and effectiveness verified as indicated above. Cells were treated with vehicle (0.1% EtOH), 100 nM Dex, or 100 nM DHT, for 1 h before the start of ChIP. ChIP experiments were done as previously indicated ^30^. Antibodies per IP: GR, 12,5 µl; p63, 1 µg; FRA1, 3 µg; GATA2, 5 µl; H3K27ac, 2 µg; AR 3 µl. Two to five IP samples were pooled for one sequencing sample. Libraries were made using NEBNext Ultra II DNA Library Prep Kit (New England BioLabs, #E7645L) according to manufacturer’s instructions. Library quality control was done with Agilent Bioanalyzer 2100 DNA 1000 Analysis kit (Agilent, #5067–1504) or NGS Fragment Kit (1-6000bp) (Agilent, DNF-473-0500). In general, two replicates were then sequenced with Illumina NextSeq 500 (75SE), Illumina NextSeq 2000 (100SE), or Illumina NovaSeq X (100PE).

For assay for transposase-accessible chromatin with sequencing (ATAC-seq), RWPE1 cells were plated on 10-cm plates (2M cells per plate). Cells were treated with vehicle (0.1% EtOH), or 100 nM Dex for 1 h before the start of nuclei isolation. ATAC-seq was performed as previously described ^31,32^. Quality analysis of ATAC-seq libraries was done using High Sensitivity DNA Analysis kit (Agilent, #5067-4626) or NGS Fragment Kit (1-6000bp) (Agilent, DNF-473-0500). Two biological replicates were sequenced using Illumina NextSeq 2000 (50PE).

### Sequential (re)ChIP-qPCR

Sequential ChIP-qPCR was performed as previously indicated ^33^. The immunocomplexes were eluted from the magnetic beads using 10 mM dithiothreitol. The subsequent ChIP was performed as the first one with antibody amount as indicated above. qPCR was performed using LightCycler 480 SYBR Green I master (Roche, #04887352001) with LightCycler 480 Instrument II (Roche). Binding was calculated with formula E−(ΔCt) × 10, where E (the efficiency of target amplification) is a coefficient of DNA amplification by one PCR cycle for a particular primer pair, and ΔCt is Ct_(ChIP template)_ – Ct_(Input)_. The used primers are listed in Supplemental Supplementary Table S1.

### ChIP-seq and ATAC-seq data analysis

ChIP-seq and ATAC-seq alignment to hg38 and read filtering was done with Bowtie2 (RRID:SCR_005476) ^34^. HOMER (4.1.1) was used for data analysis ^28^ with findPeaks used to call peaks over ChIP input sample. Replicate consensus peaks were used with RWPE1-AR data and RWPE1 p53, p63, and FRA1 ChIP-seq data. Differentially bound peaks were discovered using getDifferentialPeaksReplicates and getDifferentialPeaks. For overlapping peaks, mergePeaks using given size of the peaks were utilized. ChIP-seq peaks are presented in Supplementary Table S3. The GR ChIP-seq peaks after p63 ablation were categorized as: p63KO-lost, -gained, and -retained, based on the overlap with p63-positive control. GR peaks in Control, not present in either p63KO line were classified as p63KO-lost. GR peaks gained after p63KO, shared between p63KO lines were classified as p63KO-gained. GR peaks present in both Control and p63KO cells were classified as p63KO-retained sites. The siTP63 impacted GR and AR peak categories in RWPE1-AR cells were determined with getDifferentialPeaks command using default fold change cut off. Peaks from Dex and DexDHT treatments were combined for siTP63 differentially enriched areas to form siTP63 DN and UP categories. Downstream analyses were normalized to 10 million mapped reads. Heatmaps were generated with 10 or 20 bp bins with ± 0.5 kb or 1 kb surrounding the peak. Tag counts in box plots were normalized using log_2_ or variance stabilized and transformed (rlog) options. Box plots were drawn with Tukey method. Tag counts for scatter plots were normalized using log_2_ option. Motif enrichment was measured using findMotifsGenome.pl on default settings. Motif data is presented in Supplementary Table S4. Motif histograms were normalized to motif per bp per site and displayed ± 100-250 bp from peak center. H3K27ac histograms were normalized to tags per bp per site and displayed ± 1 kb from peak center. Motif enrichment log_2_ fold difference was calculated based on the relation of percentage target sequences with motif to the percentage of background sequences with motif. For the comparison of p63 motif and half GRE motif overlap, motif distances were called with HOMER annotatePeaks.pl and the motifs filtered based on the relative distance. The motifs were then aligned by sequence overlap. Half GRE motifs located inside p63 motifs were classified as complete overlap, while the rest of the motifs containing an overlap of two base pairs or more were classified as partial overlap. For pathway analysis, peaks were first associated with target genes using annotatePeaks.pl followed by Metascape pathway analysis like with RNA-seq data. For the prediction of transcriptional regulators from peaks, epigenetic landscape in silico deletion analysis (LISA) ^35^ from Coverage command was used with default settings and hg38 genome. Gene output from annotatePeaks.pl was utilized as the gene list, and H3K27ac BigWig files as coverage over genome.

### scRNA-seq

For single-cell RNA-seq (scRNAseq), RWPE1 cells were plated on 10-cm plates (2M cells per plate). The cells were treated with vehicle (0.1% EtOH) or 100 nM Dex for 6 hours before the start of extraction. Single cell suspensions were isolated according to published protocol and diluted to 700 cells per µl (10xGenomics, #CG00054). Library generation was performed using Single Cell 3’ Kit v3.1 (10x Genomics, #PN-1000269, #PN-1000127, #PN-1000215) according to protocol (10x Genomics, #CG000315) with target cell recovery set to 2000 cells. Library quality was done using Agilent 2100 Bioanalyzer using High Sensitivity DNA Analysis kit (Agilent, #5067-4626), and libraries were sequenced with Illumina NextSeq 2000 (100PE). The scRNA-seq data analyses were done as previously indicated ^33^. In total 4058 single cells were obtained from sequencing of EtOH (2169 cells) and Dex (1889 cells) samples. Cells with low (<750) or high (>7500) feature counts, and cells with high mitochondrial reads (>20 %) and low ribosomal reads (<5 %) were filtered out. After filtration 1468 EtOH- and 1592 Dex-treated single cells were qualified for downstream analyses. Downstream analysis was performed using Seurat 4.3.0 (RRID:SCR_007322) (Hao et al. 2021). Violin plots represent the average of the normalized expression values. Differential expression was determined with default algorithm; log_2_ fold change < 0.10, *p*-value < 0.05, min.pct =0.25). Cells with non-zero expression values were counted as expressing a transcript.

### Confocal imaging

For confocal imaging, the cells were plated on 8-well chamber slides (Ibidi GmbH, #80826) (10k cells per well). The cells were fixed with 4% paraformaldehyde (PFA) for 20 min and washed with 0.1 M phosphate buffer. The fixed cells were permeabilized for 15 minutes with 0.1% Triton X-100 in 1% BSA and stained with Phalloidin-iFluor 594 reagent (Abcam, #ab176757) for 20 min at room temperature. Nuclei were labeled with 4’,6-diamidino-2-phenylindole (DAPI) (Sigma-Aldrich, #D8417). The fluorescent images were obtained with LSM800 confocal microscope (Carl Zeiss Microscopy GmbH) using 40x objective. Image processing was performed using ZEN 3.10 software (Carl Zeiss Microscopy GmbH). For determining the cell length-to-width ratio (aspect ratio), the cell length and width in pixels were first measured with ImageJ software (National Institutes of Health, RRID:SCR_003070) and the aspect ratio was calculated by dividing the length with width (n=37-41).

### Cell proliferation assay

For automated live cell proliferation experiments, the cells were seeded onto 96-well plate (10k cells per well). At the start of the experiment (day 0), the cells were treated with vehicle (0.1% EtOH) or 100 nM Dex. For experiments involving inhibitors vehicle (DMSO 0.1%) or inhibitor 10 µM K-7174 (MedChemExpress, # HY-12743) and 1 µM GNE-7883 (MedChemExpress, # HY-147214) were added on day 0. The nuclei were stained with Incucyte Nuclight Rapid Red reagent (Essen BioSciences, #4717). The cells were monitored every day for 5 days using Incucyte S3 Live-Cell Imaging System (Sartorius). Incucyte S3 2024A software (Sartorius) was used to count the number of cells. Results were normalized to day 0 and are shown as relative percentage in cell count, representing the mean ±SD of 8-10 images per day. Box plots were drawn with Tukey method. For RNA interference, siRNA treatments were conducted as described above before plating the cells. Statistical significance was determined by Two-ANOVA with Bonferroni *post hoc* test.

### Migration assay

For automated live cell migration experiments, the cells were seeded onto 96-well plate (Sartorius Imagelock, #BA-04855) (70k cells per well). The following day, scratch wounds were generated with Incucyte Woundmaker Tool (Sartorius) and cells were washed twice with PBS. Subsequently, the cells were treated with vehicle (0.1% EtOH) or 100 nM Dex and were monitored every 3 h for total of 48 h using Incucyte S3 Live-Cell Imaging System (Sartorius). Incucyte S3 2024A software (Sartorius) was used to analyze wound healing. Results are shown as relative wound density, representing the mean ±SD of 4-5 images per time point. Box plots were drawn with Tukey method. Statistical significance was determined by Two-ANOVA with Bonferroni *post hoc* test.

### Invasion assay

For 3D invasion, the cells were plated onto 8-well chamber slides (Ibidi GmbH, #80826) (250k to 350k cells per well). The following day, Cultrex Basement Membrane Extract (R&D systems, #3432) and Type I collagen (R&D Systems, #3440) (1:1) gel mixture was added on each well and allowed to polymerize at 37°C for 2 h. Subsequently, media containing either vehicle (0.1% EtOH),100 nM Dex, or 1 µM Mifepristone (Mife) (Sigma-Aldrich, #M8046), were added on the wells and cells were allowed to invade the gel for 6 days. Fresh media was added to the slides every 48 h. The wells were fixed with 4% PFA for 2 h at room temperature and washed with 0.1 M phosphate buffer. The wells were permeabilized with 0.1% Triton-X100 in 1% BSA for 30 min and stained with Phalloidin-iFluor 594 reagent (Abcam, #ab176757) for 1 h at room temperature. Nuclei were labeled with DAPI (Sigma-Aldrich, #D8417). Cell invasion was imaged with LSM800 confocal microscope (Carl Zeiss Microscopy GmbH) using 20x objective by taking 1 µm z-intervals. Images were processed and invasion distances measured with ZEN 3.10 software (Carl Zeiss Microscopy GmbH). Box plots were drawn with Tukey method. For RNA interference, siRNA treatments were conducted as described above before plating the cells. Statistical significance was determined by One-way ANOVA with Bonferroni *post hoc* test.

### Public datasets and patient data analysis

The following publicly available sequencing datasets were used: GSE214757 for GR ChIP-seq and RNA-seq from VCaP and 22Rv1 cells ^10^, GSE204813 for RNA-seq from p63 overexpressed PC3 cells ^36^, E-MTAB-15731 for RWPE1-ARc5 ATAC-seq ^57^, GSE51497 for GR ChIP-seq in LREX cells ^14^. The data was processed as indicated above. PCa patient, survival, and single-cell expression data were obtained through Combined Transcriptome dataset of PCa Cell lines (CTPC) ^37^, Prostate cancer atlas ^38^, Xena (RRID:SCR_018938) ^39^, cBioPortal (RRID:SCR_014555) ^40^, Single Cell portal (RRID:SCR_014816) ^41^. For survival, the patients were divided based on either median or top 25% and bottom 25% quartiles. Statistical significance was calculated with log-rank test for survival.

## Results

### Chromatin binding and GR regulated transcriptome separates prostate cells from each other

For comprehensive investigation of GR action in the prostate, we examined GR-regulated transcriptome with RNA-seq and chromatin binding (cistrome) of GR with ChIP-seq from two AR-positive (VCaP, 22Rv1) ^10^ and two AR-negative (PC3, DU145) PCa cell lines (Supplementary Figure S1A-B) ^37^. Cell lines were selected based on endogenous GR expression, eliminating GR-low or -negative PCa lines such as LNCaP from the analyses. In addition, we employed immortalized normal prostate epithelial cell line RWPE1 ^42^, which express similar levels of GR as DU145 cells (Supplementary Figure S1C) ^15^, as model for non-cancerous prostate cells. Intriguingly, principal component analysis (PCA) of GR’s transcriptomes (Figure 1A) and cistromes (Figure 1B), when treated with synthetic GR agonist dexamethasone (Dex), display clear separation based on the subtype of prostate cells. VCaP and 22Rv1 formed one cluster, while AR-negative PC3 and DU145 cells formed another based on Dex induced gene expression. Moreover, GR action in RWPE1 cells differentiated from both AR-positive and -negative PCa cells. As an example, GR binding sites (GRBs) at *PER1* locus are shared between all the cell lines, while *KRT16* locus shows RWPE1-specific GRBs and *PTPRJ* locus shows primarily PCa-specific GRBs with the differences reflected at transcriptional regulation (Figure 1C, Supplemental Figure S1D). The prostate cell subtype-specific GR activity could be explained by the distinct cooperative partners of the receptor, since the transcriptional activity of GR is modulated by other TFs in PCa ^13^.

**Figure 1.**
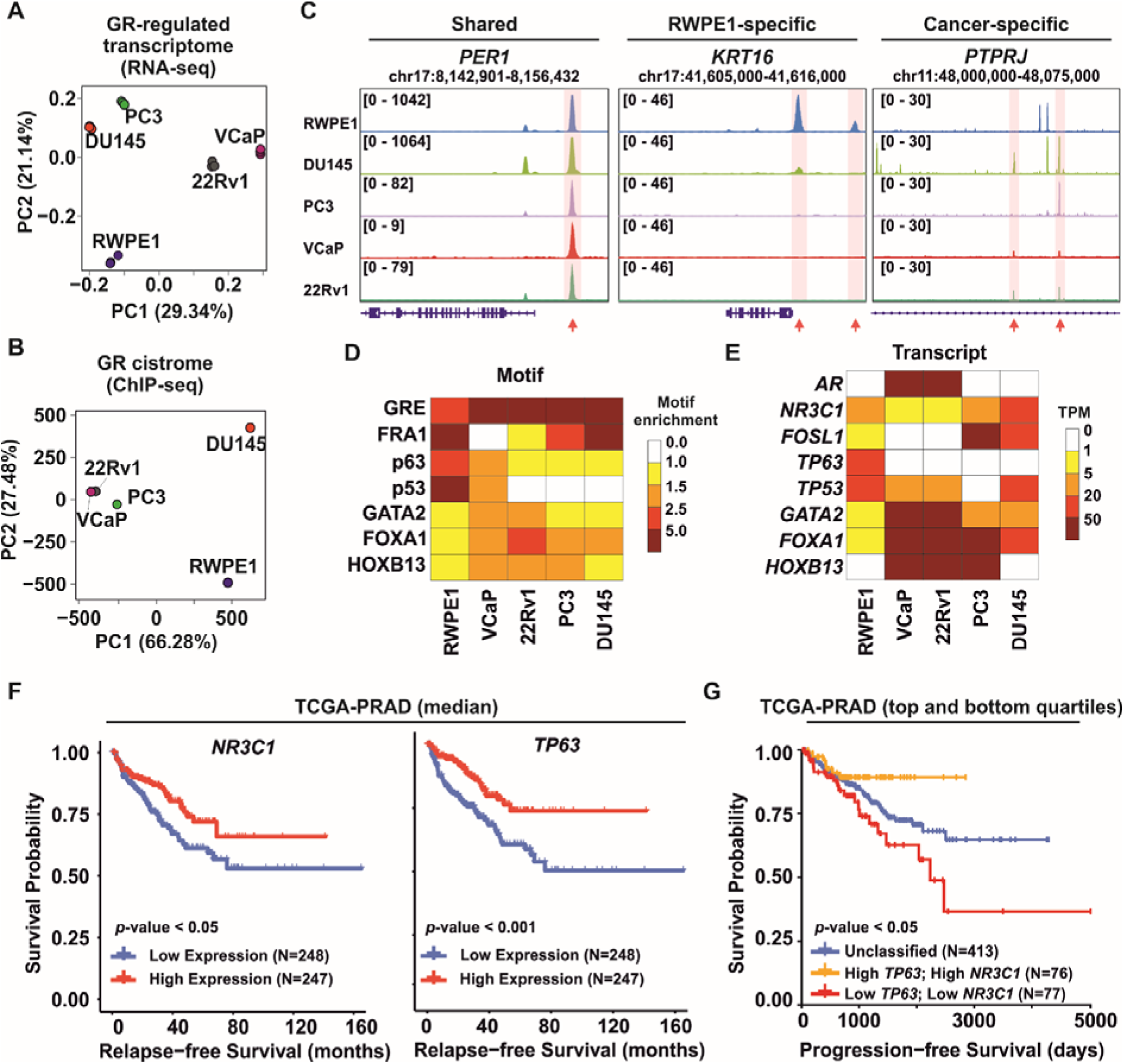
GR has distinct chromatin binding profile in normal and malignant prostate cells. (**A-B**) PCA plot of (**A**) GR-regulated transcriptome and (**B**) GR cistrome from RWPE1, VCaP, 22Rv1, PC3, and DU145 cells. (**C**) Genome browser tracks of GR ChIP-seq at *PER1*, *KRT16*, and *PTPRJ* loci. Red arrows point specific peaks for each locus. (**D-E**) Heatmap of (**D**) GRB motif enrichment and (**E**) expression of corresponding TFs. Motif enrichment values calculated as fold change over background, and expression as TPM values. (**F**) Relapse-free survival of TCGA-PRAD patients expressing either high or low levels of (left) *NR3C1* or (right) *TP63*. (**G**) Progression-free survival of TCGA-PRAD patients expressing high levels of both *TP63* and *NR3C1*, or low levels of both *TP63* and *NR3C1*. Other patients were classified as unclassified. Survival p-values calculated with log-rank test. PRAD, prostate adenocarcinoma; TCGA, the cancer genome atlas.

To investigate such TF partners, we analyzed motif enrichment of the GRBs (Supplementary Table S4). Glucocorticoid response elements (GREs) were prominently enriched in all cell lines, with PCa cells exhibiting higher enrichment than RWPE1 cells (Figure 1D, Supplemental Figure S1E). While motifs for luminal epithelial TFs, such as GATA2, FOXA1 and HOXB13 were accumulated at GRBs in VCaP and 22Rv1 cells, AP-1 motif was considerably enriched at PC3 and DU145 cells. These distinct motif enrichments are concordant with the known role of the former TFs in AR-positive PCa and the latter TF in AR-negative PCa ^6^. Most intriguingly, tumor suppressor p53-family motifs were prominently enriched only at the GRBs of RWPE1 cells. The motif enrichments matched with the TF expression levels; *GATA2*, *FOXA1*, and *HOXB13* are highly expressed in AR-positive PCa cells, *FOSL1*, subunit of AP-1, is prominently expressed in AR-negative PCa cells, and *TP53* and *TP63* are strongly expressed in RWPE1 cells (Figure 1E, Supplementary Figure S1F). These results suggest that GR’s cooperative TFs mediate its action in non-cancerous and cancerous prostate cells, likely influencing the tumor suppressive and oncogenic activities of the receptor.

To complement the RWPE1 data, we also generated GR cistrome from immortalized normal prostate epithelial cell line PNT1A ^43^. PCA of GR cistromes indicated closer association of PNT1A cells with RWPE1 than DU145 cells (Supplementary Figure S1G-H). Moreover, p53-family motifs were prominently enriched also at the GRBs of PNT1A cells (Supplementary Figure S1I), and both RWPE1 and PNT1A peaks were equally associated with p53 pathway (Supplementary Figure S1J). During the progression of PCa, the *NR3C1* expression shows minimal change, while *TP63* is the only p53-family member that is prominently decreased during cancer progression (Supplementary Figure S1K). Intriguingly, the high expression of *NR3C1* and *TP63* are associated with significantly improved relapse-free survival (Figure 1F), while no change is seen with *TP53* and *TP73* expression (Supplementary Figure S2A). Moreover, PCa patients with concurrent high expression of *NR3C1* and *TP63* have the best progression-free survival (Figure 1G). In sum, our results indicate that p53-family members, especially p63, could mediate the tumor suppressive activities of GR in the normal prostate.

### GR cistrome exhibits substantial overlap with p63 binding sites

Considering that p53-family is well-characterized for its overall significance in cancer ^44^, and normal prostate GRBs seem to be linked with p53-family members, we further explored their role in modulating GR activity. We profiled the chromatin binding of both p53 and p63 in RWPE1 cells. While the epithelial isoform DNp63α is the most abundant p63 in prostate tissue ^45^, for simplicity we will utilize p63 term throughout the text. Both TFs overlapped with GRBs, but compared to p53, p63 showed a considerable (over 70%) overlap with GRBs (Figure 2A, Supplementary Figure S2B). Moreover, p53-family motifs were enriched only at GRBs that showed p63 binding regardless of p53 occupancy (Figure 2B, Supplementary Table S4). GRBs that overlapped only with p53 showed no p53-family motifs enrichment. The lack of motif enrichment at p53 binding sites can be partly explained by the transient nature of p53 stability in non-stressed cells ^46^. Intriguingly, GRBs co-occupied with p63 showed minimal enrichment of GREs, while the center of the GRBs showed accumulation of p53 and p63 motifs (Figure 2C). Close inspection revealed the presence of a *de novo* GRE half-site motif overlapping the p63 motif (Supplementary Figure S2C). Unlike known motifs that correspond to previously characterized TF binding sequences, *de novo* motifs are sequence patterns identified directly from the analyzed genomic regions. Around 11% of the p63-GR sites had either complete (5.7%) or partial (5.6%) overlap between the half GRE and p63 motif (Supplementary Figure S2D). In addition, the p63 binding was significantly stronger in sites with either complete or partial motif overlap compared to sites without overlap (Supplementary Figure S2E). These analyses highlight that p63 is anticipated to modulate the chromatin binding of GR to its binding sites in RWPE1 cells. Based on these observations, we focused on the relationship between GR and p63.

**Figure 2.**
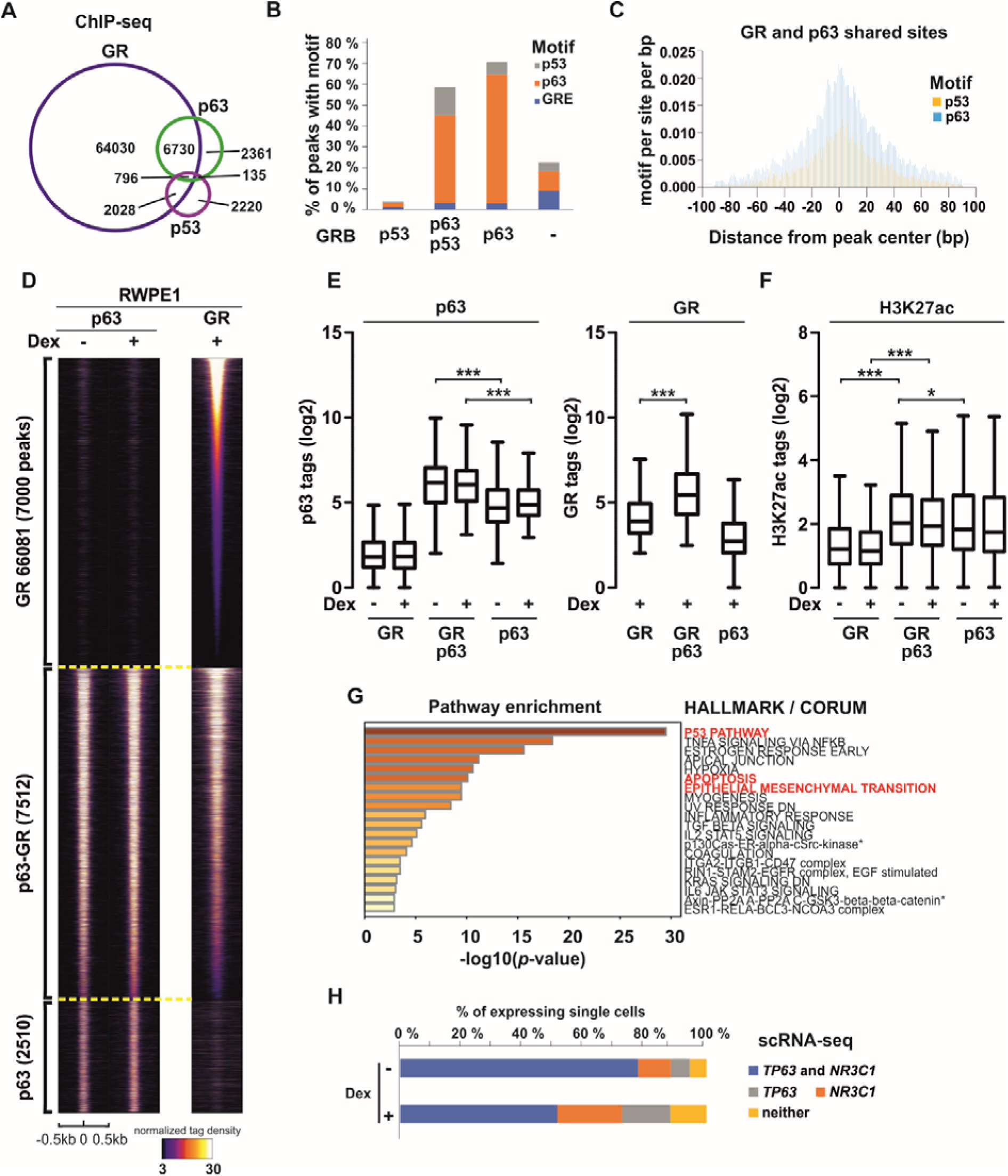
GRBs have extensive overlap with p63 in RWPE1 cells. (**A**) Overlap of GR (blue), p53 (purple), and p63 (green) ChIP-seq peaks in Dex-treated RWPE1 cells. (**B**) Percentage of p53 (grey), p63 (orange) and GRE (blue) motif enrichment at GRBs shared with only p53, with both p53 and p63, only with p63, or neither p53 nor p63. (**C**) Enrichment of p53 (yellow) and p63 (blue) motifs surrounding GR and p63 shared binding sites. (**D**) p63 and GR ChIP-seq profiles at GR, p63-GR, and p63 sites in RWPE1 cells. GR sites bind only GR, p63-GR sites bind both GR and p63, and p63 sites bind only p63. For GR sites, only 7000 randomized sites are shown in the heatmap. Each heatmap represents ± 500 bp around the center of the peak, with binding intensity (tags per bp per site) scale is noted below on a linear scale. (**E**) Log_2_ tag enrichment of (left) p63 and (right) GR at the indicated binding sites. (**F**) Log_2_ tag enrichment of H3K27ac at the indicated binding sites. (**G**) Hallmark and CORUM Gene Sets pathway analysis of p63-GR sites. Color scale represents -log_10_ *p*-value, and distinct pathways highlighted in red. Pathway names with asterisk are truncated. (**H**) Single cells expressing *NR3C1* (orange), *TP63* (grey), both (blue), or neither (yellow) in scRNA-seq data from RWPE1 cells. Expression is defined as a non-zero value. Statistical significance in box plots was calculated with One-way ANOVA with Bonferroni *post hoc* test. *, *p*< 0.05; **, *p*< 0.01; ***, *p*< 0.001.

Clustering of GR and p63 binding sites indicated that GR activation had minimal effect on the chromatin occupancy of p63 (Figure 2D-E). This suggests that p63 chromatin occupancy precedes GR activation, likely facilitating GR recruitment to these sites. Intriguingly, the binding of both p63 and GR was significantly stronger at shared sites compared to sites where they bind on their own (Figure 2E). Moreover, the active enhancer histone mark, histone 3 lysine 27 acetylation (H3K27ac), was most prominently enriched at p63-GR shared binding sites (Figure 2F). While only ∼4% of GRBs displayed at least twofold change in H3K27ac levels following GR activation, nearly half of GR-regulated genes were associated with GRBs co-occupied by p63 and GR (Supplemental Figure S2F-G). Moreover, GR-mediated changes in H3K27ac at p63-GR sites correlated with altered transcription of nearby genes (Supplementary Figure S2H). These results highlight the biological relevance of GR and p63 co-occupying sites. This is supported by pathway analyses showing a significant enrichment of p63-GR binding sites with the p53 as well as apoptosis and epithelial-mesenchymal transition (EMT) pathways (Figure 2G).

While p63 motif was specifically enriched at RWPE1 GRBs, AP-1 motifs showed enrichment at GRBs in multiple cell lines (Figure 1D). To further assess the distinctive relationship between GR and p63, we profiled chromatin binding of FRA1 in RWPE1 cells. We selected FRA1 for its prominent role in PCa ^6^. FRA1 had a prominent (over 60%) overlap with GR while its overlap with p63 was only 8% (Supplementary Figure S2I). Most p63 sites overlapped with GR without FRA1. The GRBs that harbored p63, with or without FRA1, had high enrichment whereas GRBs with FRA1 lacking p63 had low enrichment of H3K27ac (Supplementary Figure S2J). This suggests that GRBs with p63 are the most active enhancers in RWPE1 cells. Finally, single-cell RNA-seq (scRNA-seq) analyses indicate that over 75% of RWPE1 cells expressed both *NR3C1* and *TP63* in the same single cell (Figure 2H). This highlights the interconnected relationship between GR and p63. Interestingly, GR activation downregulated the transcription of both *NR3C1* and *TP63* (Supplementary Figure S2K), reducing the number of cells expressing both genes (Figure 2H). This supports the existence of negative regulatory feedback loop, as previously observed by Sevilla et al. (2024)^47^. Thus, GR and p63 crosstalk is likely a multifaceted phenomenon occurring not only at the chromatin level but also via transcriptional regulation.

Considering the physiological importance of AR in prostate epithelial cells, we examined GR and p63 action in the AR-positive RWPE1 model ^22^. Although dihydrotestosterone (DHT) modestly and context-dependently affected GR and p63 occupancy at sites identified after Dex treatment (Supplementary Figure S3A), combined treatment with Dex and DHT (DexDHT for simplicity) induced numerous additional GR-p63 binding sites (Supplementary Figure S3B). Consistently with this, GR, p63, and AR exhibited nearly complete co-occupancy on chromatin (Supplementary Figure S3C-D), supporting our previous findings that AR facilitates GR chromatin recruitment ^33^. Around three-quarters of p63-GR-AR co-occupied sites contained hormone response element (HRE) motifs, including GREs, AREs, or PREs (PR), whereas only ∼17% harbored a p63 motif (Supplementary Figure S3E). Furthermore, most co-occupied sites contained multiple high-scoring HRE motifs (Supplemental Figure S3F-G). These findings suggest that AR and GR can simultaneously occupy the same enhancer and that AR facilitates the recruitment of both GR and p63 to DHT-dependent sites enriched for highly conserved HREs. The relative scarcity of p63 motifs at these sites suggests that p63 recruitment may occur, at least in part, through protein-protein interactions rather than direct DNA binding. Together, these findings indicate that AR promotes GR and p63 chromatin occupancy which may shape transcriptional regulation in intermediate prostate epithelial cells, where AR and p63 are co-expressed.

### Loss of p63 leads to prominent changes in GR-regulated transcriptome

To investigate the degree of p63 influence on GR activity, we utilized CRISPR-Cas9 to knockout p63 from RWPE1 cells (p63KO) (see Materials and Methods for details). The utilized guide RNA (gRNA) generated a frameshift mutation resulting in premature STOP codon at exon 5 before critical amino acids for p63’s DNA binding (Supplementary Figure S4A). Two monoclonal p63KO cell lines, clone 1 (CL#1) and 2 (CL#2), were utilized in the downstream analyses (Figure 3A). RWPE1 cell line exposed to non-targeting gRNA followed by monoclonal selection was used as control (CTRL) cell line. PCA plot indicated that compared to CTRL, the transcriptome of p63KO cells, especially CL#1, moved on one axis towards the AR-negative DU145 cell transcriptome (Supplementary Figure S4B). Intriguingly, while the transcriptome of p63KO cells were close to p63-null PC3 cells, the CTRL transcriptome resembled the transcriptome of PC3 cells overexpressing p63 ^36^ (Supplementary Figure S4C). These results suggest that gain or loss of p63 leads to bidirectional transcriptomic changes. AR-positive PCa cells exhibit a luminal epithelial (LE) phenotype, while the disappearance of basal epithelial (BE) cell layers is potentially associated with PCa development ^48^. Since parental RWPE1 cells display LE-BE intermediate phenotype ^42^, and p63 is a known regulator of BE phenotype and inhibitor of EMT ^49,50^, we quantified the expression of LE, BE, and EMT marker genes in CTRL and p63KO cells.

**Figure 3.**
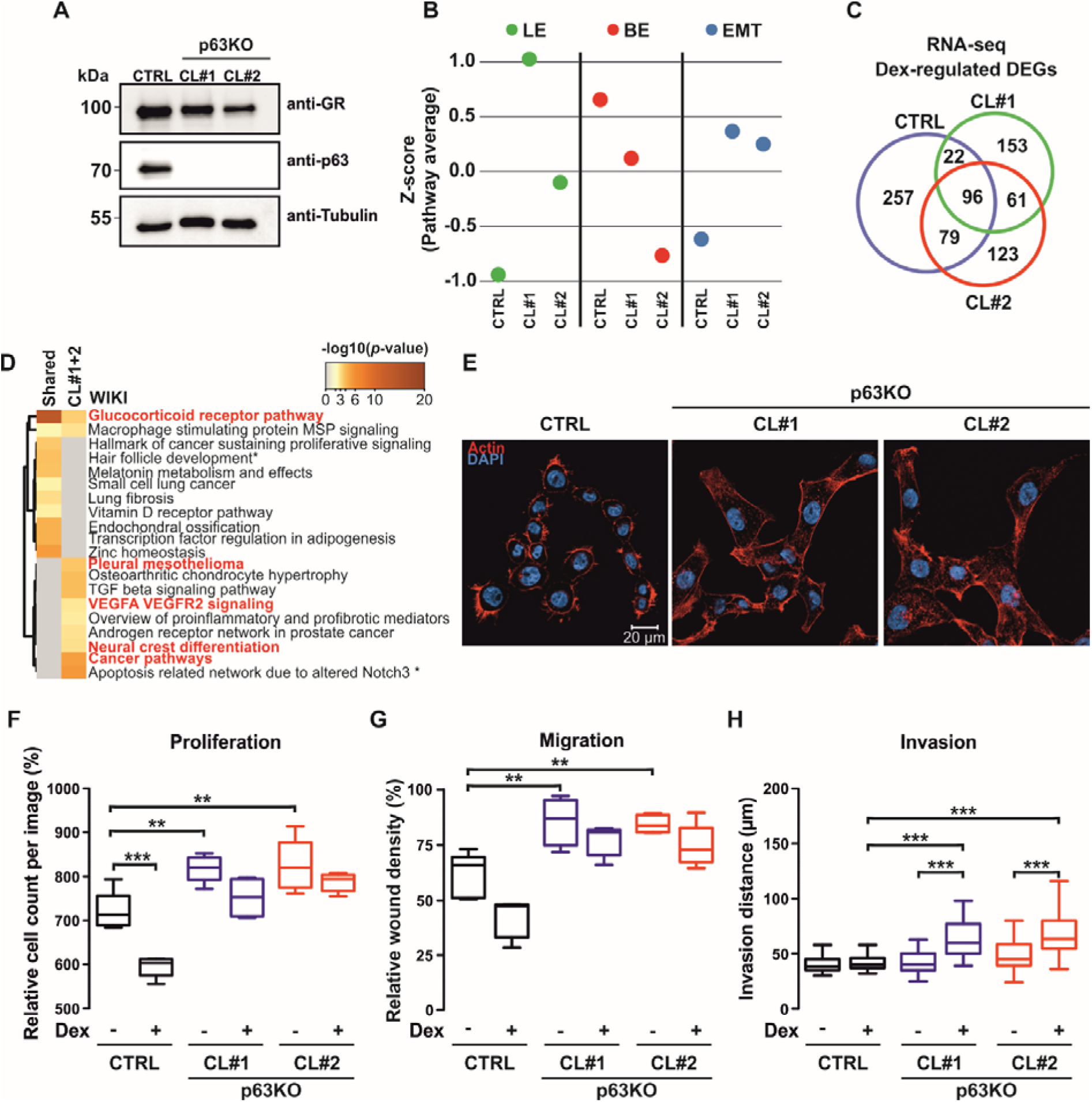
Depletion of p63 alters cell identity and EMT marker gene expression. (**A**) Immunoblotting of GR, p63, and tubulin protein levels in RWPE1 control (CTRL) and p63 depleted (p63KO) cells. Two clones (CL#1 and CL#2) of p63KO cells were utilized. (**B**) Average Z-scores for luminal epithelial (LE) (left; green), basal epithelial (BE) (middle; red), and epithelial-mesenchymal transition (EMT) (right; blue) marker gene expression in CTRL and p63KO cell lines. (**C**) Overlap of Dex-regulated DEGs in CTRL (blue), CL#1 (green) and CL#2 (red) p63KO cells. (**D**) WIKI pathway analysis of shared or CL#1 and CL#2 (CL#1+2) specific Dex-regulated DEGs. In pathway heatmaps, color scales represent -log_10_ *p*-value, and distinct pathways are highlighted in red. Pathway names with asterisk are truncated. (**E**) Immunofluorescence (IF) images of RWPE1 CTRL and p63KO cells with nuclei (DAPI; blue) and cell membrane (actin; red) staining. Scale bar, 20 μm. (**F**) Relative cell proliferation of CTRL and p63KO cells on day 5 (n=5) with indicated treatments. (**G**) Relative migration in wound healing assay of CTRL and p63KO cells on day 2 (n=8) with indicated treatments. (**H**) 3D gel invasion of CTRL and p63KO cells on day 6 with indicated treatments. Two of the highest invasion points were utilized (n=36-52). Statistical significance in box plots was calculated with One-way ANOVA with Bonferroni *post hoc* test. *, *p*< 0.05; **, *p*< 0.01; ***, *p*< 0.001.

Interestingly, the loss of p63 led to phenotypic switch wherein p63KO cells expressed more LE and less BE marker genes compared to CTRL (Figure 3B, Supplementary Figure S4D-E). Despite the increase in LE marker genes, we observed no *AR* or *KLK3* expression upon p63KO. In parallel with LE marker gene induction, genes associated with EMT were prominently increased upon p63KO (Figure 3B, Supplementary Figure S4F). This was corroborated at pathway level with differentially regulated genes (DEGs) in p63KO displaying a prominent enrichment of EMT pathway (Supplementary Figure S4G-H). Moreover, other cancer progression related pathways, such as VEGFA VEGFR2 signaling and embryonic stem cell pluripotency, were enriched with p63KO DEGs. Focusing on Dex-regulated DEGs, the loss of p63 lead to a clear transcriptomic change where several genes gained or lost their Dex-mediated regulation (Figure 3C, Supplementary Figure S5A). Intriguingly, while GR pathway was enriched with Dex-regulated genes in both CTRL and p63KO cells, only the latter showed enrichment of PCa malignancy related pathways, such as VEGFA VEGFR2 signaling, neural crest differentiation, and cancer pathways (Figure 3D, Supplementary Figure S5A). Consistently with the findings, several EMT markers, including TWIST1 and NCAD, were Dex-regulated in p63KO cells (Supplementary Figure S5A-B). Notably, TWIST1 and NCAD protein levels reached their highest levels after 5 days of Dex treatment (Supplementary Figure S5B), suggesting that prolonged GR activation promotes accumulation of oncogenic mediators. In p63KO cells, long-term Dex exposure had only a minor impact on GR protein levels abundance. In contrast, both GR and p63 protein levels were prominently reduced in CTRL cells. These results imply that p63 loss alters GR to have a more oncogenic potential in prostate cells.

### Absence of p63 leads to pronounced changes in cell morphology, proliferation, migration, and invasion

Given the p63-depletion induced expression of LE and EMT marker genes and GR’s prominent role in PCa cell survival ^15^, we sought to investigate whether these changes corresponded with enhanced oncogenic potential. In agreement with phenotypic transcriptome changes, p63KO cells showed altered morphology with round shape converted to a more jagged and elongated appearance (Figure 3E, Supplementary Figure S5C). The opposite effect has been seen with PC3 cells where overexpression of p63 converts cells’ elongated shape to a rounder appearance ^51^. The p63KO-mediated change in morphology was associated with higher baseline proliferation rates and attenuation of the anti-proliferative effects of Dex (Figure 3F, Supplementary Figure S6A-B). Moreover, a similar significant increase in migration and suppression of the anti-migrative effects of Dex was observed in p63KO compared to CTRL cells (Figure 3G, Supplementary Figure S6C-E). The concordant effect on migration is seen with overexpression of p63 in PC3 cells ^51^. Thus, increased proliferation and migration are driven by the loss of p63 while GR’s effect on these changes is diminished.

Since EMT pathway was prominently enriched with p63KO DEGs, including Dex-regulated genes, we measured the 3D invasiveness of the cells (see Materials and Methods for details). While proliferation and migration displayed significant changes, the cells’ invasiveness showed only slight change upon p63KO (Figure 3H, Supplementary Figure S6F). However, Dex treatment led to significant increase in p63KO cells’ invasiveness, which was not observed in CTRL cells (Figure 3H, Supplementary Figure S6F). Dex-induced invasion in p63KO cells was abolished by the GR antagonist mifepristone (Supplementary Figure S6G).

These findings indicate p63 loss enables GR-dependent invasive programs, coinciding with the accumulation of EMT markers during prolonged Dex exposure (Figure 3C-D, Supplementary Figure S5A-B). Therefore, the loss of p63 shifts GR behavior from tumor suppressive towards more oncogenic activity.

### Reprogramming GR cistrome upon loss of p63

Due to the clear Dex-induced changes in the cells’ invasiveness, we profiled GR cistrome from p63KO cells. Comparison of GR cistromes indicated that p63KO led to both loss and gain of GRBs (Supplementary Figure S7A). Similar to transcriptomes (Supplementary Figure S4B), PCA plot showed that the GR cistromes of p63KO cells, especially CL#1, advanced towards the GRB profile of PCa cell lines (Supplementary Figure S7B). As an example of GRBs, *MAOA* and *GNAO1* loci showed loss of p63 co-occupying GRBs, while *GCNT1* locus displayed gain of GRB (Figure 4A). Moreover, these changes were correlated at gene regulatory level, as *MAOA* and *GNAO1* lost while *GCNT1*, associated with extracapsular PCa occurrence ^52^, gained Dex-induction in p63KO cells (Figure 4B). Several GR-regulated genes exhibited altered basal transcription levels, which may partially explain the increased baseline proliferation observed in these cells (Supplementary figure S7C). To investigate what drives GRBs changes, we performed motif enrichment analyses from the p63KO GRBs (Supplementary Table S4). In accordance with the depletion of p63, the p53-family motifs displayed the most prominent loss at GRBs (Figure 4C, Supplementary Figure S7D). At transcript level only *TP63* showed significant reduction, indicating that the loss of p63 drives the changes. While GR itself, *i.e.*, GRE enrichment and *NR3C1* transcript levels, remained largely unaffected, *FOSL1* transcripts were significantly increased in p63KO cells. However, GRBs showed no prominent difference in AP-1 motif enrichment. In comparison to the mixed AP-1 differences, p63KO cells exhibited a distinct increase in GATA-family motif enrichment and elevated *GATA2* transcript levels (Figure 4C).

**Figure 4.**
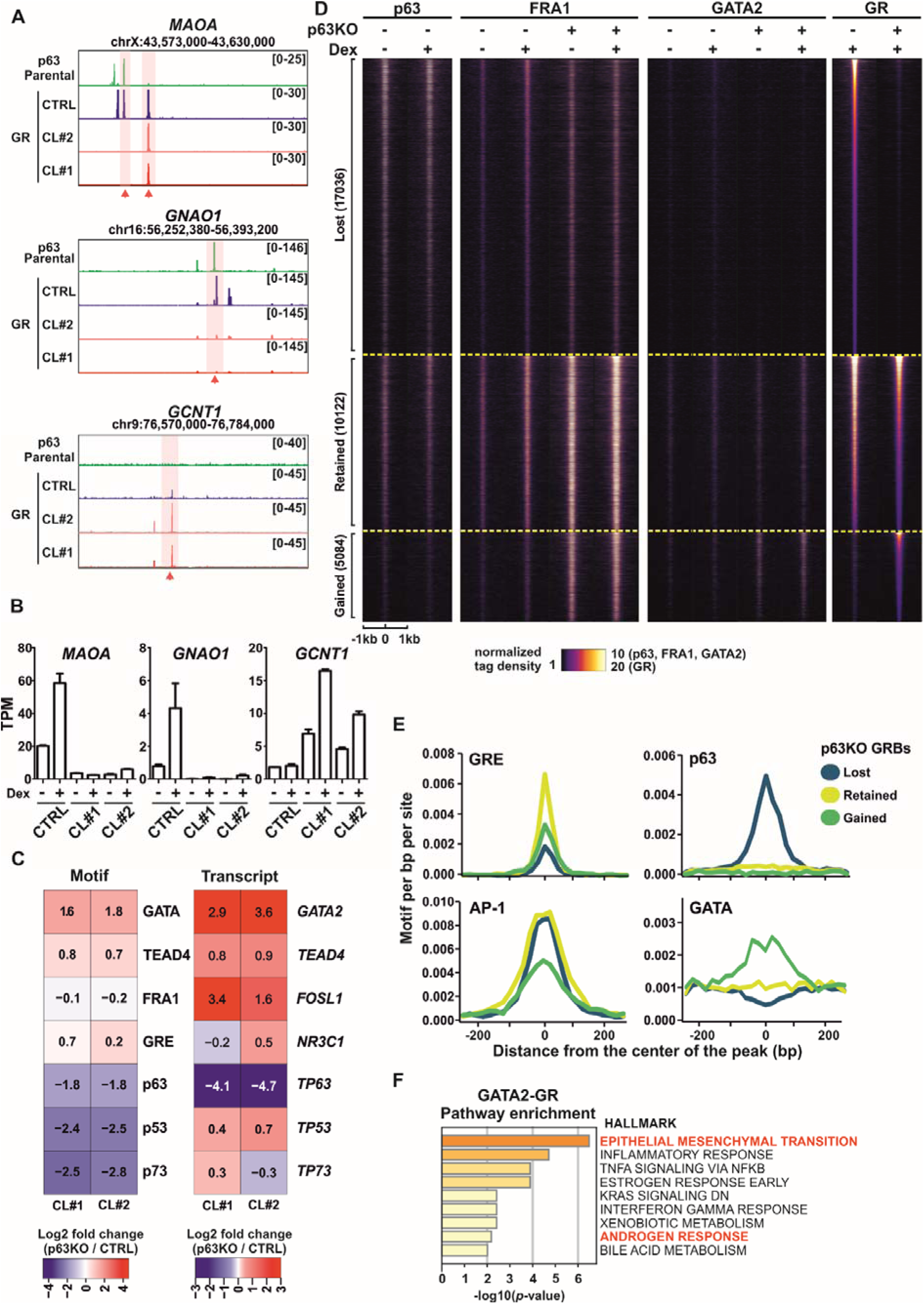
Depletion of p63 modulates the GR cistrome. (**A**) Genome browser tracks of p63 ChIP-seq from parental RWPE1 cells, and GR ChIP-seq from CTRL and p63KO cells at *MAOA*, *GNAO1*, and *GCNT1* loci. Red arrows point specific peaks for each locus. (**B**) RNA-seq TPM values of *MAOA*, *GNAO1*, and *GCNT1* from EtOH- and Dex-treated CTRL and p63KO cells. Bar graphs represent mean ±SD, n=2. (**C**) Heatmap of GRB motif enrichment and corresponding TF expression change in p63KO cells. Motif enrichment and expression values calculated as log2 fold change over CTRL cells. (**D**) ChIP-seq profiles of p63, FRA1, GATA2, and GR from p63-intact (p63KO -) and p63KO (p63KO +) cells at p63KO-lost, -retained, and -gained GRBs. Each heatmap represents ± 1 kb around the center of the peak, and binding intensity (tags per bp per site) is noted below on a linear scale. (**E**) Motif density of GRE, p63, AP-1, and GATA at indicated p63KO GRBs. Histograms represent ± 240 bp around the center of the peak, and enrichment intensity represents motif per bp per site. (**F**) Hallmark Gene Sets pathway analysis of GATA2-GR sites. Color scale represents -log_10_ *p*-value, and distinct pathways highlighted in red.

To better examine what mediates the GRB changes upon p63KO, we divided GR cistromes to p63KO-lost, - retained, and -gained GRBs (see Materials and Methods for details) (Figure 4D, Supplementary Figure S7E). The lost and retained GRBs were enriched at the p53 pathway, whereas only gained GRBs showed prominent enrichment of androgen response pathway (Supplementary Figure S7F). This suggests that both gained GRBs and Dex-regulated genes in p63KO cells (Figure 3D), mimic the PCa cells’ regulatory profiles. In addition, the chromatin accessibility and H3K27ac enrichment were significantly decreased at lost GRBs whereas they were significantly increased at gained GRBs upon p63KO (Supplementary Figure S8A-B).

These results highlight that p63 mediates the enhancer activity of lost GRBs, while other TFs likely have a reciprocal effect at gained GRBs. To identify such TFs, we performed motif analyses. While GREs were enriched at all GRBs, with the clearest enrichment at retained GRBs, p63 motif was enriched only at lost GRBs (Figure 4E). The latter was corroborated with p63 ChIP-seq data from parental cells, indicating that lost GRBs were the most p63 occupied sites (Figure 4D, Supplementary Figure S7E). Moreover, p63 silencing in RWPE1-AR cells reduced GR occupancy at nearly half of the p63KO-lost sites (Supplementary Figure S8C). This suggests that disruption of p63-dependent GR recruitment is a major contributor to the loss of these binding events, rather than secondary effects arising from clonal adaptation of p63KO cells.

Furthermore, our reChIP assays indicated co-occupancy of p63 and GR on EGFL7 enhancer locus (Supplementary Figure S8D), which corroborates previous research in keratinocytes ^47^. Even though AP-1 is a known mediator of GR action ^53^ and AP-1 motifs showed modest enrichment at gained GRBs, the most prevalent enrichment of AP-1 motifs were observed at lost and retained GRBs. FRA1 ChIP-seq data from the p63KO cells indicated higher occupancy on retained GRBs and moderate FRA1 binding on gained sites (Figure 4D, Supplementary Figure S7E). This suggests that although *FOSL1* transcript levels are increased, FRA1 alone is not the primary mediator of GR binding to gained GRBs in p63KO cells.

Intriguingly, gained GRBs presented prominent enrichment of GATA-family motifs that were not observed at lost or retained GRBs (Figure 4E). Of the GATA-family members, only *GATA2* and *GATA3* showed a significant increase upon p63KO (Supplementary Figure S9A), and the promoters of *GATA2* and *GATA3* displayed significantly increased chromatin accessibility and H3K27ac enrichment in p63KO compared to CTRL cells (Supplementary Figure S9B). Interestingly, GATA2 has a prominent role in PCa ^54^, and its expression is high in early-stages of PCa (Supplementary Figure S9C). Moreover, LE marker expression was increased in p63KO cells and GATA2 is associated with luminal differentiation. Thus, we focused on the role of GATA2. First, we confirmed the prominent increase of GATA2 protein levels in p63KO cells (Supplementary Figure S9D), followed by GATA2 ChIP-seq analyses from CTRL and p63KO cells. In agreement with the increased protein levels, more GATA2 occupied sites were observed in p63KO compared to CTRL cells (Supplementary Figure S9E). More importantly, GATA2 binds to the gained GRBs in p63KO cells being absent from the lost GRBs, and – based on our reChIP results —co-occupies GR bound regions (Figure 4D, Supplementary Figure S7E, Supplementary Figure S8D). Furthermore, the GATA2-GR binding sites were associated with EMT and androgen response pathways (Figure 4F). Our results suggest that both p63 and GATA2 function as pioneer factors, as previously indicated ^55,56^, mediating GR’s occupancy at distinct sites in asynchronous fashion. The loss of p63 leads to increased GATA2 levels resulting in the reshaping of GR chromatin occupancy from p63 to GATA2 binding sites.

Consistent with these findings, siRNA-mediated depletion of p63 (siTP63) in RWPE1-AR cells (Supplementary Figure S10A) substantially reprogrammed both GR and AR chromatin occupancy profiles (Supplementary Figure S10B). Loss of p63 resulted in both reduced (siTP63 DN) and newly acquired (siTP63 UP) GR and AR binding sites. Interestingly, siTP63 UP sites were predominantly detected under the DexDHT treatment, suggesting that their formation is largely AR dependent (Supplementary Figure S10C). No such pattern was observed at siTP63 DN sites. Motif analysis revealed the expected enrichment of p63 motifs at siTP63 DN regions, while siTP63 UP regions were strongly enriched for HRE motifs (Supplementary Figure S10D). Consistent with p63KO data, p63 depletion increased the prevalence of GATA2 motifs and elevated GATA2 protein levels (Supplementary Figure S10D-E). Genes associated with siTP63 UP sites were enriched in pathways similar to those regulated by Dex in p63KO cells, including VEGFA VEGFR2 signaling, neural crest cell migration during development, and cancer pathways (Supplementary Figure S10F). Integration of published RWPE1-AR ATAC-seq data ^57^, indicated reduced chromatin accessibility at siTP63 DN sites, whereas DHT treatment increased accessibility siTP63 UP sites. Collectively, our results highlight a complex interplay between p63 and steroid receptors in prostate cells, in which AR and p63 reciprocally shape each other’s chromatin occupancy and cooperate in directing GR recruitment to their regulatory sites.

### GR’s harmful effects in p63-null cells can be partially mitigated with GATA2 depletion

Our results highlight that GR’s chromatin action is reprogrammed through the crosstalk with p63 and GATA2 with the former mediating GR’s tumor suppressive-like effects while the latter conveys GR’s oncogenic activities. Our analysis of GR cistromes supported this by indicating a higher overlap or GRBs between PCa cell lines and p63KO cells compared to p63-intact CTRL cells (Supplementary Figure S11A). Intriguingly, scRNA-seq data from human and mouse prostate tissue as well as from PCa patients shows the general mutually exclusive cell type-specific expression of *TP63* and *GATA2*, wherein *TP63* is expressed mainly in basal cells while *GATA2* is expressed in luminal and PCa cells (Figure 5A). In comparison, *NR3C1* is expressed in most cells, suggesting that other TFs, such as p63 and GATA2, mediates its cell type-specific action. At the level of survival, the high expression of *GATA2* is associated with decreased relapse-free survival (Supplementary Figure S11B), which is opposite to the *NR3C1* and *TP63* expression (Figure 1F).

**Figure 5.**
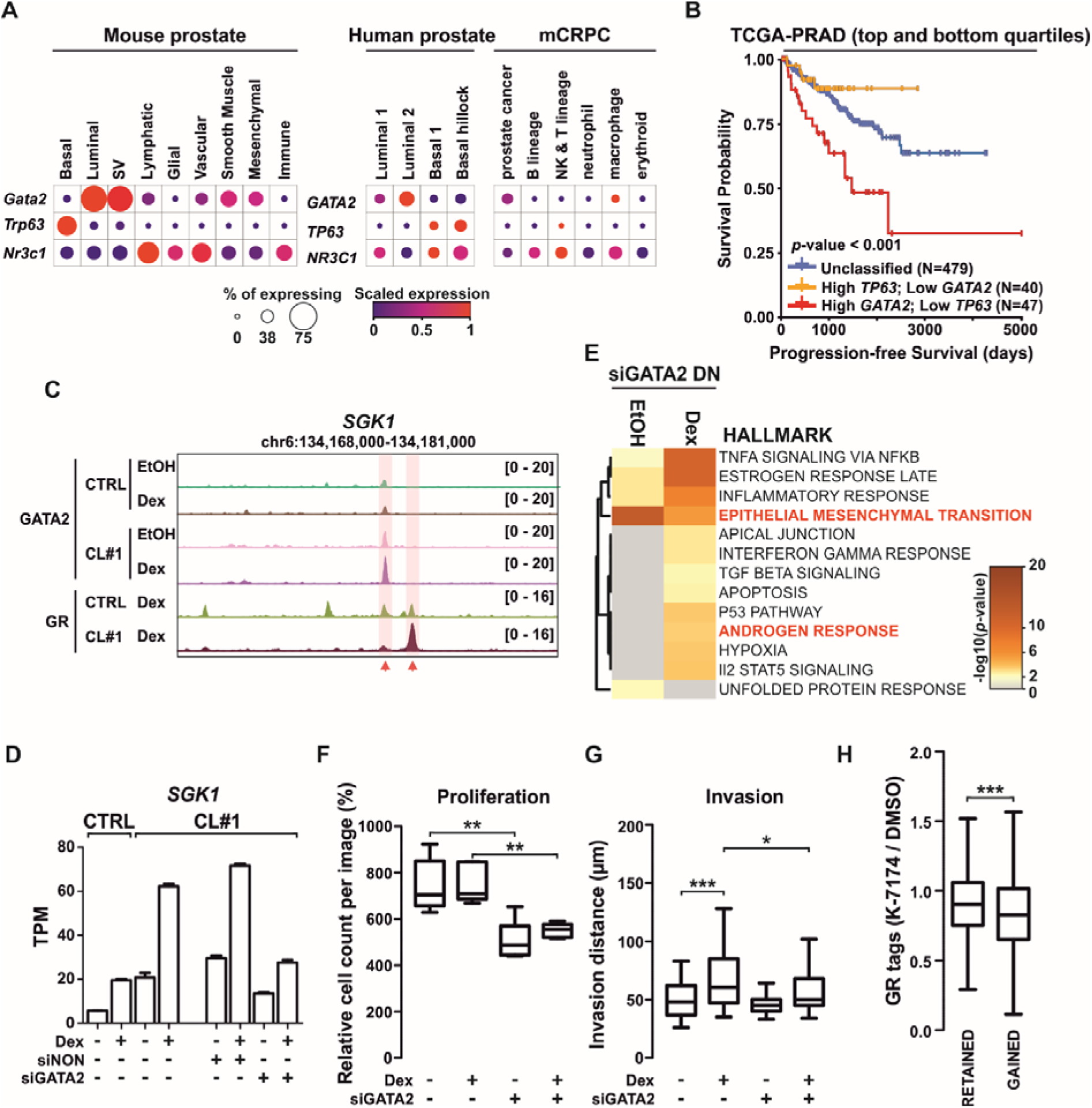
GATA2 promotes GR-induced invasion. (**A**) Single cell expression of *Gata2*, *Trp63* and *Nr3c1* in mouse prostate, and *GATA2*, *TP63* and *NR3C1* in human prostate and metastatic (m)CRPC samples. Data obtained from Single Cell Portal, and intensity of the color depicts the scaled expression, and size of the circle depicts the percentage of cells that express the gene. (**B**) Progression-free survival of TCGA-PRAD patients expressing high levels of both *TP63* and *NR3C1*, or low levels of both *TP63* and *NR3C1*. Other patients were classified as unclassified. (**C**) Genome browser track of GR and GATA2 ChIP-seq from CTRL and p63KO cells at *SGK1* locus. (**D**) RNA-seq TPM values of *SGK1* from EtOH- and Dex-treated CTRL and p63KO cells, and siNON- and siGATA2-treated p63KO cells. Bar graphs represent mean ±SD, n=2. (**E**) Hallmark Gene Sets pathway analysis of siGATA2 downregulated (DN) genes where Dex-regulation is lost or without Dex-regulation. Color scale represents -log_10_ *p*-value, and distinct pathways highlighted in red. (**F**) Relative cell proliferation of p63KO cells on day 4 (n=5) with indicated treatments. (**G**) 3D gel invasion of p63KO cells on day 6 with indicated treatments. Two of the highest invasion points were utilized (n=50-54). (**H**) Box plot depicting indicated ratio of GR ChIP-seq tags in p63KO-retained and -gained GRBs when treated with DMSO or K-7174 in p63KO cells. Statistical significance in box plots was calculated with One-way ANOVA with Bonferroni *post hoc* test. *, *p*< 0.05; **, *p*< 0.01; ***, *p*< 0.001. PRAD, prostate adenocarcinoma; TCGA, the cancer genome atlas.

More importantly, PCa patients with the high *TP63* and low *GATA2* expression have the best while high *GATA2* and low *TP63* expression have the worst progression-free survival (Figure 5B). This indicates that the differential expression of *TP63* and *GATA2* has clinical significance possibly through altering GR activity.

To further evaluate GATA2’s contribution, we depleted it from the p63KO cells using siRNA knockdown (Supplementary Figure S11C), and analyzed transcriptomic changes with RNA-seq. Depletion of GATA2 altered the expression of numerous transcripts including *AREG*, *ITGB1*, and *SGK1* (Supplementary Figure S11D). As an example, *SGK1* is a known GR target gene that shows increased GATA2 and GR binding upon p63KO (Figure 5C), which lost its p63KO-potentiated Dex-induction when GATA2 was depleted (Figure 5D). Pathway analyses indicated that siGATA2 downregulated genes were enriched at EMT pathways (Figure 5E, Supplementary Figure S11E). Similar enrichment was observed with Dex-regulated genes that showed significant attenuation of Dex response upon siGATA2 treatment. Importantly, only siGATA2-attenuated Dex-regulated genes were enriched at androgen response pathway (Figure 5E). Thus, while GATA2 regulates EMT in Dex-dependent and -independent manner, GR together with GATA2 is responsible for the regulation of androgen response pathway (Figure 3D, Figure 4F).

Since EMT and androgen response pathway were impacted by siGATA2, we analyzed the proliferation and invasion of the cells after GATA2 depletion. The proliferation of p63KO cells was significantly inhibited with siGATA2 treatment (Figure 5F, Supplementary Figure S11F). However, while anti-proliferative effects of Dex were prominent in p63-intact cells (Figure 3F), GATA2 depletion did not have any effect on the Dex responsiveness of p63KO cells (Figure 5F, Supplementary Figure S11F). In comparison, GATA2 depletion significantly impaired the GR-induced invasiveness of p63KO cells (Figure 5G). The GR-mediated invasive effects were also attenuated using K-7174 (Supplementary Figure S11G), a small-molecule GATA inhibitor^58,59^. Moreover, our GR ChIP-seq results indicate that K-7174 treatment reduced GR binding at gained GRBs in p63KO cells (Figure 5H). This suggests that the crosstalk between GATA2 and GR regulates the GR binding and expression of EMT pathway manifesting in increased cell invasion.

### GATA2 and FRA1 orchestrate oncogenic GR activity

Since depletion of GATA2 did not completely remove all the GR effects induced by p63KO, we investigated other potential contributing TFs using epigenetic landscape in silico deletion analysis (LISA) ^35^. GRBs with GATA2 were associated with GATA-family members, as expected, whereas AP-1 subunit FRA2 (encoded by *FOSL2*) was highly associated with GRBs without GATA2 (Supplementary Figure S12A). Since the expression of another AP-1 subunit, FRA1 (encoded by *FOSL1*), was significantly increased in p63KO compared to CTRL cells, we examined its overlap with p63KO gained GRBs (Figure 4C). FRA1 binding was associated with higher chromatin accessibility at GR and GATA2 occupied sites, before and after GR chromatin binding (Supplementary Figure S12B). In concordance, FRA1 has been implicated in maintaining chromatin accessibility in AR-negative advanced PCa ^6^, highlighting its potential role as a pioneer factor. Supporting this notion, GR activation had only a limited effect on FRA1 as well as GATA2 chromatin occupancy (Supplementary Figure S12C), suggesting that these TFs establish permissive chromatin landscape for GR binding. Furthermore, FRA1 occupancy mitigated the K 7174 -induced reduction in GR binding at GATA2 occupied GRBs, whereas GRBs bound only by FRA1 exhibited only a modest decrease in GR binding (Figure 6A, Supplementary Figure S12D). This suggests that FRA1 modulates chromatin accessibility and GR binding in p63KO cells, both independently and in cooperation with GATA2.

**Figure 6.**
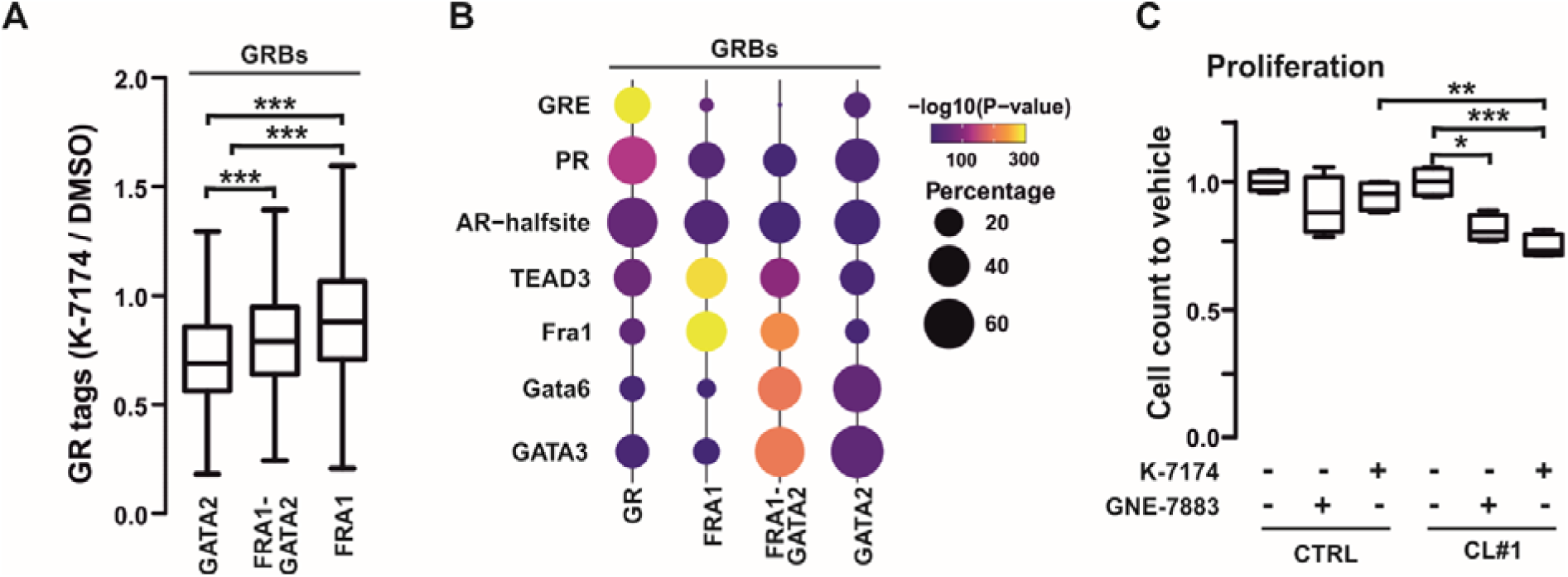
GATA2 and FRA1 co-orchestrate GR binding on gained sites. (**A**) Box plot depicting indicated ratio of GR ChIP-seq tags in DMSO or K-7174-treated p63KO cells at p63KO-gained GRBs, categorized by overlap with GATA2 and FRA1 ChIP-seq peaks. (**B**) Motif enrichment of p63KO-gained GRBs, categorized by overlap with GATA2 and FRA1 ChIP-seq peaks. GR-category indicates sites with no overlap with GATA2 or FRA1. The color represents -log_10_(p-value) of motif enrichment, and the dot size the percentage of peaks with the motif. (**C**) Relative cell proliferation of CTRL and p63KO cells on day 4 (n=4) following treatment with vehicle, GNE-7883, or K-7174. Values are normalized to the vehicle-treated average of each cell line. Statistical significance was determined with One-way ANOVA with Bonferroni *post hoc* test. *, *p*< 0.05; **, *p*< 0.01; ***, *p*< 0.001.

In support, p63KO-gained GRBs that lacked GATA2 binding contained less GREs and more FRA1 and TEAD motifs (Figure 6B, Supplementary Figure S12E). In addition, TEAD family expression, in particular *TEAD1* and *TEAD4,* exhibited increased transcription in p63KO cells (Supplementary Figure S12F). The increase of both FRA1 and TEAD is particularly intriguing, since FRA1-TEAD signaling pathways have been associated with AR-negative advanced PCa ^6^. To assess the effects of FRA1-TEAD signaling on proliferation we utilized GNE-7883, an allosteric pan-TEAD inhibitor, that prevents coactivators YAP/TAZ from binding the TEAD protein ^60^. While TEAD inhibition significantly reduced the proliferation, GATA inhibitor K-7174 displayed stronger anti-proliferative effects in p63KO cells (Figure 6C). Interestingly, GATA2 and FRA1 exhibit regulatory crosstalk, as GATA2 binds *FOSL1* promoter and *FOSL1* expression was significantly decreased upon siGATA2 treatment (Supplementary Figure S12G). This suggests that GATA2 positively regulates *FOSL1* transcription, driving the FRA1 induced oncogenic effects. For example, the expression of EMT marker *VIM* was downregulated upon GATA2 depletion in p63KO cells possibly stemming from lowered levels of FRA1 expression (Supplementary Figure S12H). This indicates that GATA2 and FRA1 cooperate on several transcriptional levels inducing both GR-dependent and -independent oncogenic effects.

Our results suggest that GR’s tumor suppressive and oncogenic action in prostate cells is largely dependent on its crosstalk partner. With p63, GR regulates basal and p53 pathway genes restricting the proliferation and migration of cells. Upon loss of p63, GR switches crosstalk partners to GATA2 and FRA1. With GATA2 and FRA1, GR regulates luminal and EMT pathway genes inducing the invasiveness of cells. Thus, GR’s expression can remain unchanged during the initiation and progression of PCa, but its action can be drastically altered by other TFs mediating its chromatin activities.

## Discussion

Glucocorticoids are among the most widely prescribed drugs globally ^61^ and are extensively used to mitigate treatment-related side effects in cancer patients ^12^. In PCa, supplemental glucocorticoids are particularly important to counteract the suppression of serum cortisol levels and prevent a compensatory rise in adrenocorticotropic hormone caused by abiraterone treatment. However, the role of the GR in PCa is complex. Studies have demonstrated that GR expression is lowered in several cancers, including prostate, and that GR possesses both pro-and anti-tumorigenic effects ^10,14–16,62^, which underscores the clinical importance of understanding when GR functions as a tumor suppressor versus as an oncogene ^13^. Despite this significance, GR’s context-dependent activity and the underlying molecular mediators have remained elusive. Here, we identify an oncogenic molecular switch that governs GR’s transition from a tumor-suppressive to an oncogenic state.

Our genome-wide analyses revealed that GR’s chromatin binding and transcriptional responses differ markedly between PCa cells and immortalized prostate epithelial cells. In PCa cells, GR activity was closely associated with the TFs FOXA1, GATA2, and AP-1, all well-established oncogenic drivers in PCa ^6^ with FOXA1 and GATA2 specifically mediating GR’s pro-tumorigenic functions ^10,63,64^. In contrast, our data indicated that in immortalized prostate cells, GR predominantly associated with the p53-family member p63. Our findings demonstrate that p63 plays a central role in directing GR’s anti-tumorigenic activity. Given that GR expression remains relatively constant across normal and malignant prostate tissues ^65,66^, and that GR function shifts during PCa progression ^13^, it is likely that differential TF crosstalk, rather than GR abundance, underlies its context-dependent activity.

PCa typically exhibits a loss of BE cells, while its origin is most commonly traced to LE cells ^48,67^. However, BE cells can also serve as a source of PCa following their differentiation into an LE-like state ^68^. Our investigation reveals that p63-null prostate cells display a diminished BE phenotype and an enhanced LE-like identity, consistent with the well-established role of p63 as a key regulator of BE cell fate ^49,50^. In addition to this shift in epithelial identity, p63-null cells exhibit notable morphological changes and elevated expression of EMT markers, supporting previous findings that p63 suppresses EMT in the prostate ^50,69^. Intriguingly, despite the upregulation of EMT markers, GR activation was required to drive the invasive behavior of p63-null cells.

In p63-intact prostate cells, GR regulated p53-associated gene expression programs through active enhancers co-occupied by both GR and p63. At these shared regulatory sites, GR suppressed cell proliferation and migration. Patient data supported these tumor-suppressive effects, showing the most favorable survival outcomes in individuals with high expression of both GR and p63. However, the protective role of GR was markedly altered upon p63 depletion. In p63-null prostate cells, GR’s chromatin binding profile was reprogrammed, losing occupancy at p63-bound enhancers and gaining access to GATA2-bound enhancers. From these newly acquired sites, GR activated oncogenic transcriptional programs, including GR-induced accumulation of EMT marker genes, leading to diminished anti-proliferative effects and enhanced cell invasiveness. These molecular changes were mirrored in patient survival data, which revealed the best outcomes in tumors with high p63 and low GATA2 expression, and the poorest outcomes in those with high GATA2 and low p63 expression. Nonetheless it is to be noted, that higher p63 expression may be associated with lower ratio of basal cell displacement thus skewing the survival data. To offset potential sampling bias in PCa survival data, approaches such as single-cell transcriptomics integrated with clinical outcomes could help disentangle biological phenomena from technical effects. Furthermore, lineage-tracing models could shed light on whether aberrant p63 loss induces differentiation into luminal-like cells capable of tumor initiation.

Although GR and p63 have not previously been linked in the prostate, their cooperative roles have been documented in other disease contexts. GR and p63 have been shown to co-occupy chromatin in basal-like pancreatic ductal adenocarcinoma cells ^70^. In keratinocytes, GR and p63 exhibit substantial overlap in chromatin binding, and physically interact at shared regulatory sites ^47^. Notably, GR loss in this context leads to atopic dermatitis, driven by a p63-mediated increase in inflammatory signaling, suggesting that GR counteracts the detrimental effects of p63 in skin. Similarly, in the prostate, our findings indicate that GR and p63 cooperate to suppress tumorigenic processes, highlighting a beneficial partnership in disease attenuation. However, this cooperative effect is likely not universally protective. While p63 restricts EMT in PCa, it has been shown to promote EMT in other malignancies, such as squamous cell carcinoma ^45,71,72^. These findings underscore the context- and disease-specific nature of GR–p63 crosstalk, which may yield either protective or pathogenic outcomes depending on cellular and molecular environments. Because p63 overexpression shifts PCa cell migration ^51^ and transcriptome ^36^ toward a more normal prostate phenotype, it is plausible that GR-mediated transcriptional regulation is similarly affected. Further studies examining gene expression and chromatin occupancy will be required to test this possibility.

Given the central role of AR signaling in prostate differentiation and cancer, we also employed an AR-positive prostate model that co-express AR, GR and p63 ^22^. Interestingly, both p63 and AR appear to modulate chromatin accessibility and to facilitate the recruitment of GR, as well as each other, to their respective binding sites. The ability of AR to influence GR occupancy is consistent with our previous findings demonstrating that, in PCa cells, AR expands GR cistrome and thereby promotes a distinct transcriptional outcome ^33^. In RWPE1-AR cells, however, the impact of AR appeared even more pronounced, extending to the recruitment of p63. Notably, AR activation reduced chromatin accessibility at p63-dependent GRBs independently of GR activation. This might reflect the ability of AR to redistribute p63 toward HRE-enriched regulatory regions, in line with the previous observations that AR activation reduces chromatin accessibility at stemness-related loci ^57^. The capacity of AR to modulate p63 activity has intriguing implications for prostate epithelial differentiation, as this interplay may contribute to the transition of intermediate lineage cells toward a luminally differentiated state. Although loss of p63 resulted in increased GATA2 levels in AR-positive prostate cells, most changes in GR occupancy required concomitant AR activation, suggesting that GATA2 plays a relatively minor role in GR recruitment in the presence of active AR. Further studies are needed to determine if aberrant loss of p63 promotes in harmful reprogramming of GR activity in AR-positive context.

GATA2 is a well-established oncogenic factor in PCa, known to promote disease progression through both AR-dependent and -independent mechanisms ^9,54,55,73^. Most notably, GATA2 enhances AR signaling by modulating AR expression and facilitating its chromatin binding ^9,55^ and contributes to antiandrogen resistance by regulating GR expression ^64^. Our findings extend GATA2’s oncogenic role by demonstrating its involvement in reprogramming GR activity following p63 depletion-induced increase of GATA2. In p63-null prostate cells, GR’s acquired regulation of EMT and androgen response pathways was abrogated upon GATA2 silencing. Additionally, GATA2 inhibition reduced GR binding to sites associated with GATA2. Furthermore, GATA2 depletion and chemical inhibition significantly reduced GR-driven cell invasion. These results highlight GATA2 as a critical mediator of GR’s oncogenic reprogramming.

Although the interplay between AP-1 and GATA family members has not been extensively characterized in PCa, their cooperative activity at shared enhancers has been documented in other biological contexts. For example, GATA2 and AP-1 can jointly reprogram human fibroblasts into hematopoietic progenitor cells ^74^, and in breast cancer, GATA3 collaborates with AP-1 to establish functional enhancers ^75^. Notably, p63 depletion leads to increased expression of AP-1 subunits, such as *FOSL1*, at levels comparable to GATA2. Furthermore, depletion of GATA2 reduced the expression of *FOSL1* implicating GATA2 as a modulator of AP-1. In p63-null prostate cells, GRBs with or without GATA2 were enriched with FRA1 binding sites, which also buffered the reduction of GR binding to GATA2 occupied sites when treated with GATA inhibitor. This suggests that AP-1 acts jointly with GATA2 to modulate GR activity on a number of sites while also having independent effects. Thus, AP-1 may also contribute to GR’s oncogenic activity in the absence of p63. Indeed, AP-1 is a well-established mediator of GR action ^53^ and together with TEAD proteins it has been implicated as a driver of stem cell-like PCa ^6^. Relatedly, p63KO induced a moderate upregulation of TEAD family proteins. Inhibiting TEAD by GNE-7883 ^60^, a small-molecule inhibitor, specifically reduced cell proliferation in p63KO cells, indicating an increased reliance on TEAD signaling. GNE-7883 has been shown to suppress chromatin accessibility on AP-1 motifs in malignant mesothelioma cells, highlighting the interconnectedness of TEAD1 and AP-1 function ^76^. The involvement of AP-1 provides a plausible explanation for the persistence of some oncogenic effects of GR in p63-null cells even after GATA2 depletion, highlighting a broader network of TF crosstalk driving GR-mediated tumor progression.

Our findings suggest that TF crosstalk, governs the transformation of GR activity from tumor-suppressive to oncogenic in prostate cells. This mechanism highlights a broader principle of steroid receptor plasticity. A similar, though mechanistically distinct, phenomenon has been observed in breast cancer, where the tumor-suppressive functions of the AR can be reprogrammed into oncogenic activity via the JAK/STAT pathway ^21^. Notably, our study demonstrates that such reprogramming of a steroid receptor can occur even in non-cancerous prostate cells. Furthermore, our results point to GATA2 and AP-1 as potential therapeutic targets to mitigate the detrimental effects of glucocorticoids in PCa. Given that GATA2 also regulates *NR3C1* expression in antiandrogen-treated PCa cells ^64^, targeting GATA2 may be an effective strategy to suppress GR-driven oncogenic activity across diverse PCa contexts. Small-molecule GATA2 inhibitor K-7174 ^58,64^ holds promise as its effects were specifically observed in p63-null cells. Moreover, additional GATA inhibitors such as repurposed vasodilator Dilazep ^77^, offer promising avenues for intervention, although these compounds have yet to be evaluated in clinical trials. AP-1 presents an interesting therapeutic target; however, the functional complexity of AP-1 subunits complicates drug development, and no AP-1 inhibitors have been clinically approved.

## Data availability

The generated ChIP-seq, ATAC-seq, RNA-seq, and scRNA-seq datasets have been submitted to the NCBI Gene Expression Omnibus database (http://www.ncbi.nlm.nih.gov/geo/). Accession code: GSE301472.

## Acknowledgements

We thank Siri Mäki-Ikola, Merja Räsänen, Saara Pirinen, and Kirsi Kainulainen for technical assistance. The EMBL GeneCore team (Heidelberg, Germany) is greatly acknowledged for deep sequencing services. This work was carried out with the support of UEF Single Cell Genomics Core and UEF Cell and Tissue Imaging Unit, University of Eastern Finland, Finland, Biocenter Kuopio and EuroBioimaging Finland. The computational analyses were performed on servers provided by UEF Bioinformatics Center, University of Eastern Finland, Finland.

## Author contributions

**Johannes Hiltunen**: Conceptualization; Formal Analysis; Investigation; Visualization; Writing – original draft; Writing – review & editing. **Niina Aaltonen**: Formal Analysis; Investigation; Visualization; Writing – review & editing. **Heini Sohlberg**: Investigation; Writing – review & editing. **Laura Kemppi**: Investigation; Writing – review & editing. **Konsta Kukkonen**: Resources; Writing – review & editing. **Tapio Visakorpi**: Resources; Writing – review & editing. **Ville Paakinaho**: Conceptualization; Formal Analysis; Funding acquisition; Investigation; Project administration; Supervision; Visualization; Writing – original draft; Writing – review & editing.

## References

1. Bray F, Laversanne M, Sung H, Ferlay J, Siegel RL, Soerjomataram I, et al. Global cancer statistics 2022: GLOBOCAN estimates of incidence and mortality worldwide for 36 cancers in 185 countries. CA Cancer J Clin. 2024;74(3):229–63. doi:10.3322/caac.21834 PubMed PMID: 38572751.

2. Wang G, Zhao D, Spring DJ, DePinho RA. Genetics and biology of prostate cancer. Genes Dev. 2018 Sep 1;32(17–18):1105–40. doi:10.1101/gad.315739.118 PubMed PMID: 30181359; PubMed Central PMCID: PMC6120714.

3. Feng Q, He B. Androgen Receptor Signaling in the Development of Castration-Resistant Prostate Cancer. Front Oncol. 2019;9:858. doi:10.3389/fonc.2019.00858 PubMed PMID: 31552182; PubMed Central PMCID: PMC6738163.

4. Paakinaho V, Palvimo JJ. Genome-wide crosstalk between steroid receptors in breast and prostate cancers. Endocr Relat Cancer. 2021 Sep 1;28(9):R231–50.

5. Burris TP, de Vera IMS, Cote I, Flaveny CA, Wanninayake US, Chatterjee A, et al. International Union of Basic and Clinical Pharmacology CXIII: Nuclear Receptor Superfamily-Update 2023. Pharmacol Rev. 2023 Nov;75(6):1233–318. doi:10.1124/pharmrev.121.000436 PubMed PMID: 37586884; PubMed Central PMCID: PMC10595025.

6. Tang F, Xu D, Wang S, Wong CK, Martinez-Fundichely A, Lee CJ, et al. Chromatin profiles classify castration-resistant prostate cancers suggesting therapeutic targets. Science. 2022 May 27;376(6596):eabe1505. doi:10.1126/science.abe1505 PubMed PMID: 35617398; PubMed Central PMCID: PMC9299269.

7. Mu P, Zhang Z, Benelli M, Karthaus WR, Hoover E, Chen CC, et al. SOX2 promotes lineage plasticity and antiandrogen resistance in TP53- and RB1-deficient prostate cancer. Science. 2017 Jan 6;355(6320):84–8. doi:10.1126/science.aah4307 PubMed PMID: 28059768; PubMed Central PMCID: PMC5247742.

8. Linder S, Hoogstraat M, Stelloo S, Eickhoff N, Schuurman K, de Barros H, et al. Drug-Induced Epigenomic Plasticity Reprograms Circadian Rhythm Regulation to Drive Prostate Cancer toward Androgen Independence. Cancer Discov. 2022 Sep 2;12(9):2074–97. doi:10.1158/2159-8290.CD-21-0576 PubMed PMID: 35754340; PubMed Central PMCID: PMC7613567.

9. Zhou T, Yu C, Han Y, He B, Feng Q. GATA2 up-regulation restores androgen receptor chromatin association and advances darolutamide resistance in prostate cancer. Genes Dis. 2025 Jul;12(4):101508. doi:10.1016/j.gendis.2024.101508 PubMed PMID: 40201140; PubMed Central PMCID: PMC11978331.

10. Helminen L, Huttunen J, Tulonen M, Aaltonen N, Niskanen EA, Palvimo JJ, et al. Chromatin accessibility and pioneer factor FOXA1 restrict glucocorticoid receptor action in prostate cancer. Nucleic Acids Res. 2024 Jan 25;52(2):625–42. doi:10.1093/nar/gkad1126

11. Williams A, Gutgesell L, de Wet L, Selman P, Dey A, Avineni M, et al. SOX2 expression in prostate cancer drives resistance to nuclear hormone receptor signaling inhibition through the WEE1/CDK1 signaling axis. Cancer Lett. 2023 Jul 1;565:216209. doi:10.1016/j.canlet.2023.216209 PubMed PMID: 37169162.

12. Kalfeist L, Galland L, Ledys F, Ghiringhelli F, Limagne E, Ladoire S. Impact of Glucocorticoid Use in Oncology in the Immunotherapy Era. Cells. 2022 Feb 22;11(5):770. doi:10.3390/cells11050770 PubMed PMID: 35269392; PubMed Central PMCID: PMC8909189.

13. Hiltunen J, Helminen L, Paakinaho V. Glucocorticoid receptor action in prostate cancer: the role of transcription factor crosstalk. Front Endocrinol. 2024 Jul 4;15:1437179. doi:10.3389/fendo.2024.1437179 PubMed PMID: 39027480; PubMed Central PMCID: PMC11254642.

14. Arora VK, Schenkein E, Murali R, Subudhi SK, Wongvipat J, Balbas MD, et al. Glucocorticoid receptor confers resistance to antiandrogens by bypassing androgen receptor blockade. Cell. 2013 Dec 5;155(6):1309–22. doi:10.1016/j.cell.2013.11.012 PubMed PMID: 24315100; PubMed Central PMCID: PMC3932525.

15. Puhr M, Hoefer J, Eigentler A, Ploner C, Handle F, Schaefer G, et al. The Glucocorticoid Receptor Is a Key Player for Prostate Cancer Cell Survival and a Target for Improved Antiandrogen Therapy. Clin Cancer Res Off J Am Assoc Cancer Res. 2018 Feb 15;24(4):927–38. doi:10.1158/1078-0432.CCR-17-0989 PubMed PMID: 29158269.

16. Yemelyanov A, Czwornog J, Chebotaev D, Karseladze A, Kulevitch E, Yang X, et al. Tumor suppressor activity of glucocorticoid receptor in the prostate. Oncogene. 2007 Mar 22;26(13):1885–96. doi:10.1038/sj.onc.1209991 PubMed PMID: 17016446.

17. Rogerson C, Britton E, Withey S, Hanley N, Ang YS, Sharrocks AD. Identification of a primitive intestinal transcription factor network shared between esophageal adenocarcinoma and its precancerous precursor state. Genome Res. 2019 May;29(5):723–36. doi:10.1101/gr.243345.118 PubMed PMID: 30962179; PubMed Central PMCID: PMC6499311.

18. Parreno V, Loubiere V, Schuettengruber B, Fritsch L, Rawal CC, Erokhin M, et al. Transient loss of Polycomb components induces an epigenetic cancer fate. Nature. 2024 May;629(8012):688–96. doi:10.1038/s41586-024-07328-w PubMed PMID: 38658752; PubMed Central PMCID: PMC11096130.

19. Hickey TE, Selth LA, Chia KM, Laven-Law G, Milioli HH, Roden D, et al. The androgen receptor is a tumor suppressor in estrogen receptor-positive breast cancer. Nat Med. 2021 Feb;27(2):310–20. doi:10.1038/s41591-020-01168-7 PubMed PMID: 33462444.

20. Prekovic S, Chalkiadakis T, Roest M, Roden D, Lutz C, Schuurman K, et al. Luminal breast cancer identity is determined by loss of glucocorticoid receptor activity. EMBO Mol Med. 2023 Dec 7;15(12):e17737. doi:10.15252/emmm.202317737 PubMed PMID: 37902007; PubMed Central PMCID: PMC10701603.

21. Asemota S, Effah W, Holt J, Johnson D, Cripe L, Ponnusamy S, et al. A molecular switch from tumor suppressor to oncogene in ER+ve breast cancer: Role of androgen receptor, JAK-STAT, and lineage plasticity. Proc Natl Acad Sci U S A. 2024 Oct;121(40):e2406837121. doi:10.1073/pnas.2406837121 PubMed PMID: 39312663; PubMed Central PMCID: PMC11459127.

22. Kukkonen K, Autio-Kimura B, Rauhala H, Kesseli J, Nykter M, Latonen L, et al. Nonmalignant AR-positive prostate epithelial cells and cancer cells respond differently to androgen. Endocr Relat Cancer. 2022 Dec 1;29(12):717–33. doi:10.1530/ERC-22-0108 PubMed PMID: 36219867; PubMed Central PMCID: PMC9644224.

23. Chen C, Gorlatova N, Kelman Z, Herzberg O. Structures of p63 DNA binding domain in complexes with half-site and with spacer-containing full response elements. Proc Natl Acad Sci. 2011 Apr 19;108(16):6456–61. doi:10.1073/pnas.1013657108

24. Karvonen U, Kallio PJ, Jänne OA, Palvimo JJ. Interaction of androgen receptors with androgen response element in intact cells. Roles of amino-and carboxyl-terminal regions and the ligand. J Biol Chem. 1997 Jun 20;272(25):15973–9. doi:10.1074/jbc.272.25.15973 PubMed PMID: 9188499.

25. Launonen KM, Paakinaho V, Sigismondo G, Malinen M, Sironen R, Hartikainen JM, et al. Chromatin-directed proteomics-identified network of endogenous androgen receptor in prostate cancer cells. Oncogene. 2021 Jul;40(27):4567–79. doi:10.1038/s41388-021-01887-2 PubMed PMID: 34127815; PubMed Central PMCID: PMC8266679.

26. Dobin A, Davis CA, Schlesinger F, Drenkow J, Zaleski C, Jha S, et al. STAR: ultrafast universal RNA-seq aligner. Bioinforma Oxf Engl. 2013 Jan 1;29(1):15–21. doi:10.1093/bioinformatics/bts635 PubMed PMID: 23104886; PubMed Central PMCID: PMC3530905.

27. Love MI, Huber W, Anders S. Moderated estimation of fold change and dispersion for RNA-seq data with DESeq2. Genome Biol. 2014;15(12):550. doi:10.1186/s13059-014-0550-8 PubMed PMID: 25516281; PubMed Central PMCID: PMC4302049.

28. Heinz S, Benner C, Spann N, Bertolino E, Lin YC, Laslo P, et al. Simple combinations of lineage-determining transcription factors prime cis-regulatory elements required for macrophage and B cell identities. Mol Cell. 2010 May 28;38(4):576–89. doi:10.1016/j.molcel.2010.05.004 PubMed PMID: 20513432; PubMed Central PMCID: PMC2898526.

29. Zhou Y, Zhou B, Pache L, Chang M, Khodabakhshi AH, Tanaseichuk O, et al. Metascape provides a biologist-oriented resource for the analysis of systems-level datasets. Nat Commun. 2019 Apr 3;10(1):1523. doi:10.1038/s41467-019-09234-6 PubMed PMID: 30944313; PubMed Central PMCID: PMC6447622.

30. Paakinaho V, Kaikkonen S, Makkonen H, Benes V, Palvimo JJ. SUMOylation regulates the chromatin occupancy and anti-proliferative gene programs of glucocorticoid receptor. Nucleic Acids Res. 2014 Feb;42(3):1575–92. doi:10.1093/nar/gkt1033 PubMed PMID: 24194604; PubMed Central PMCID: PMC3919585.

31. Buenrostro JD, Wu B, Chang HY, Greenleaf WJ. ATAC-seq: A Method for Assaying Chromatin Accessibility Genome-Wide. Curr Protoc Mol Biol. 2015 Jan 5;109:21.29.1-21.29.9. doi:10.1002/0471142727.mb2129s109 PubMed PMID: 25559105; PubMed Central PMCID: PMC4374986.

32. Paakinaho V, Lempiäinen JK, Sigismondo G, Niskanen EA, Malinen M, Jääskeläinen T, et al. SUMOylation regulates the protein network and chromatin accessibility at glucocorticoid receptor-binding sites. Nucleic Acids Res. 2021 Feb 26;49(4):1951–71. doi:10.1093/nar/gkab032 PubMed PMID: 33524141; PubMed Central PMCID: PMC7913686.

33. Hiltunen J, Helminen L, Aaltonen N, Launonen KM, Laakso H, Malinen M, et al. Androgen receptor-mediated assisted loading of the glucocorticoid receptor modulates transcriptional responses in prostate cancer cells. Genome Res. 2025 Jun 2;gr.280224.124. doi:10.1101/gr.280224.124 PubMed PMID: 40456604.

34. Langmead B, Salzberg SL. Fast gapped-read alignment with Bowtie 2. Nat Methods. 2012 Mar 4;9(4):357–9. doi:10.1038/nmeth.1923 PubMed PMID: 22388286; PubMed Central PMCID: PMC3322381.

35. Qin Q, Fan J, Zheng R, Wan C, Mei S, Wu Q, et al. Lisa: inferring transcriptional regulators through integrative modeling of public chromatin accessibility and ChIP-seq data. Genome Biol. 2020 Feb 7;21(1):32. doi:10.1186/s13059-020-1934-6 PubMed PMID: 32033573; PubMed Central PMCID: PMC7007693.

36. Sultanov R, Mulyukina A, Zubkova O, Fedoseeva A, Bogomazova A, Klimina K, et al. TP63–TRIM29 axis regulates enhancer methylation and chromosomal instability in prostate cancer. Epigenetics Chromatin. 2024 Mar 14;17:6. doi:10.1186/s13072-024-00529-7 PubMed PMID: 38481282; PubMed Central PMCID: PMC10938740.

37. Cheng S, Yu X. CTPC, a combined transcriptome data set of human prostate cancer cell lines. The Prostate. 2023;83(2):158–61. doi:10.1002/pros.24448

38. Bolis M, Bossi D, Vallerga A, Ceserani V, Cavalli M, Impellizzieri D, et al. Dynamic prostate cancer transcriptome analysis delineates the trajectory to disease progression. Nat Commun. 2021 Dec 2;12(1):7033. doi:10.1038/s41467-021-26840-5 PubMed PMID: 34857732; PubMed Central PMCID: PMC8640014.

39. Goldman MJ, Craft B, Hastie M, Repečka K, McDade F, Kamath A, et al. Visualizing and interpreting cancer genomics data via the Xena platform. Nat Biotechnol. 2020 Jun;38(6):675–8. doi:10.1038/s41587-020-0546-8 PubMed PMID: 32444850; PubMed Central PMCID: PMC7386072.

40. Cerami E, Gao J, Dogrusoz U, Gross BE, Sumer SO, Aksoy BA, et al. The cBio cancer genomics portal: an open platform for exploring multidimensional cancer genomics data. Cancer Discov. 2012 May;2(5):401–4. doi:10.1158/2159-8290.CD-12-0095 PubMed PMID: 22588877; PubMed Central PMCID: PMC3956037.

41. Tarhan L, Bistline J, Chang J, Galloway B, Hanna E, Weitz E. Single Cell Portal: an interactive home for single-cell genomics data [Internet]. bioRxiv; 2023 [cited 2025 Jun 17]. p. 2023.07.13.548886. Available from: https://www.biorxiv.org/content/10.1101/2023.07.13.548886v1 doi:10.1101/2023.07.13.548886

42. Bello D, Webber MM, Kleinman HK, Wartinger DD, Rhim JS. Androgen responsive adult human prostatic epithelial cell lines immortalized by human papillomavirus 18. Carcinogenesis. 1997 Jun;18(6):1215–23. doi:10.1093/carcin/18.6.1215 PubMed PMID: 9214605.

43. Degeorges A, Hoffschir F, Cussenot O, Gauville C, Le Duc A, Dutrillaux B, et al. Recurrent cytogenetic alterations of prostate carcinoma and amplification of c-myc or epidermal growth factor receptor in subclones of immortalized PNT1 human prostate epithelial cell line. Int J Cancer. 1995 Sep 15;62(6):724–31. doi:10.1002/ijc.2910620613 PubMed PMID: 7558421.

44. Pflaum J, Schlosser S, Müller M. p53 Family and Cellular Stress Responses in Cancer. Front Oncol. 2014 Oct 21;4. doi:10.3389/fonc.2014.00285

45. Li Y, Giovannini S, Wang T, Fang J, Li P, Shao C, et al. p63: a crucial player in epithelial stemness regulation. Oncogene. 2023 Nov;42(46):3371–84. doi:10.1038/s41388-023-02859-4

46. Tsvetkov P, Reuven N, Shaul Y. Ubiquitin-independent p53 proteasomal degradation. Cell Death Differ. 2010 Jan;17(1):103–8. doi:10.1038/cdd.2009.67

47. Sevilla LM, Pons-Alonso O, Gallego A, Azkargorta M, Elortza F, Pérez P. Glucocorticoid receptor controls atopic dermatitis inflammation via functional interactions with P63 and autocrine signaling in epidermal keratinocytes. Cell Death Dis. 2024 Jul 28;15(7):535. doi:10.1038/s41419-024-06926-w PubMed PMID: 39069531; PubMed Central PMCID: PMC11284228.

48. Brawer MK, Peehl DM, Stamey TA, Bostwick DG. Keratin immunoreactivity in the benign and neoplastic human prostate. Cancer Res. 1985 Aug;45(8):3663–7. PubMed PMID: 2410099.

49. Bergholz J, Xiao ZX. Role of p63 in Development, Tumorigenesis and Cancer Progression. Cancer Microenviron. 2012 Jul 31;5(3):311. doi:10.1007/s12307-012-0116-9 PubMed PMID: 22847008.

50. Olsen JR, Oyan AM, Rostad K, Hellem MR, Liu J, Li L, et al. p63 attenuates epithelial to mesenchymal potential in an experimental prostate cell model. PloS One. 2013;8(5):e62547. doi:10.1371/journal.pone.0062547 PubMed PMID: 23658742; PubMed Central PMCID: PMC3641034.

51. Tucci P, Agostini M, Grespi F, Markert EK, Terrinoni A, Vousden KH, et al. Loss of p63 and its microRNA-205 target results in enhanced cell migration and metastasis in prostate cancer. Proc Natl Acad Sci. 2012 Sep 18;109(38):15312–7. doi:10.1073/pnas.1110977109

52. Kojima Y, Yoneyama T, Hatakeyama S, Mikami J, Sato T, Mori K, et al. Detection of Core2 β-1,6-N-Acetylglucosaminyltransferase in Post-Digital Rectal Examination Urine Is a Reliable Indicator for Extracapsular Extension of Prostate Cancer. PLoS ONE. 2015 Sep 21;10(9):e0138520. doi:10.1371/journal.pone.0138520 PubMed PMID: 26390303; PubMed Central PMCID: PMC4577128.

53. Biddie SC, John S, Sabo PJ, Thurman RE, Johnson TA, Schiltz RL, et al. Transcription factor AP1 potentiates chromatin accessibility and glucocorticoid receptor binding. Mol Cell. 2011 Jul 8;43(1):145–55. doi:10.1016/j.molcel.2011.06.016 PubMed PMID: 21726817; PubMed Central PMCID: PMC3138120.

54. Rodriguez-Bravo V, Carceles-Cordon M, Hoshida Y, Cordon-Cardo C, Galsky MD, Domingo-Domenech J. The role of GATA2 in lethal prostate cancer aggressiveness. Nat Rev Urol. 2017 Jan;14(1):38–48. doi:10.1038/nrurol.2016.225 PubMed PMID: 27872477; PubMed Central PMCID: PMC5489122.

55. Wu D, Sunkel B, Chen Z, Liu X, Ye Z, Li Q, et al. Three-tiered role of the pioneer factor GATA2 in promoting androgen-dependent gene expression in prostate cancer. Nucleic Acids Res. 2014 Apr 1;42(6):3607–22. doi:10.1093/nar/gkt1382

56. Yu X, Singh PK, Tabrejee S, Sinha S, Buck MJ. ΔNp63 is a pioneer factor that binds inaccessible chromatin and elicits chromatin remodeling. Epigenetics Chromatin. 2021 Apr 17;14(1):20. doi:10.1186/s13072-021-00394-8

57. Kukkonen K, Afyounian E, Rauhala H, Nykter M, Urbanucci A, Latonen L, et al. Androgen receptor drives divergent chromatin accessibility programs in benign and malignant prostate epithelial cells. Nucleic Acids Res. 2026 Sep 23;54(17):gkag891. doi:10.1093/nar/gkag891

58. Umetani M, Nakao H, Doi T, Iwasaki A, Ohtaka M, Nagoya T, et al. A novel cell adhesion inhibitor, K-7174, reduces the endothelial VCAM-1 induction by inflammatory cytokines, acting through the regulation of GATA. Biochem Biophys Res Commun. 2000 Jun 7;272(2):370–4. doi:10.1006/bbrc.2000.2784 PubMed PMID: 10833420.

59. Yuan F, Hankey W, Wu D, Wang H, Somarelli J, Armstrong AJ, et al. Molecular determinants for enzalutamide-induced transcription in prostate cancer. Nucleic Acids Res. 2019 Nov 4;47(19):10104–14. doi:10.1093/nar/gkz790

60. Hagenbeek TJ, Zbieg JR, Hafner M, Mroue R, Lacap JA, Sodir NM, et al. An allosteric pan-TEAD inhibitor blocks oncogenic YAP/TAZ signaling and overcomes KRAS G12C inhibitor resistance. Nat Cancer. 2023 Jun;4(6):812–28. doi:10.1038/s43018-023-00577-0

61. Pofi R, Caratti G, Ray DW, Tomlinson JW. Treating the Side Effects of Exogenous Glucocorticoids; Can We Separate the Good From the Bad? Endocr Rev. 2023 Nov 9;44(6):975–1011. doi:10.1210/endrev/bnad016 PubMed PMID: 37253115; PubMed Central PMCID: PMC10638606.

62. Matthews LC, Berry AA, Morgan DJ, Poolman TM, Bauer K, Kramer F, et al. Glucocorticoid receptor regulates accurate chromosome segregation and is associated with malignancy. Proc Natl Acad Sci. 2015 Apr 28;112(17):5479–84. doi:10.1073/pnas.1411356112

63. Huttunen J, Aaltonen N, Helminen L, Rilla K, Paakinaho V. EP300/CREBBP acetyltransferase inhibition limits steroid receptor and FOXA1 signaling in prostate cancer cells. Cell Mol Life Sci CMLS. 2024 Apr 2;81(1):160. doi:10.1007/s00018-024-05209-z PubMed PMID: 38564048; PubMed Central PMCID: PMC10987371.

64. Yuan F, Hankey W, Wu D, Wang H, Somarelli J, Armstrong AJ, et al. Molecular determinants for enzalutamide-induced transcription in prostate cancer. Nucleic Acids Res. 2019 Nov 4;47(19):10104–14. doi:10.1093/nar/gkz790 PubMed PMID: 31501863; PubMed Central PMCID: PMC6821169.

65. Karthaus WR, Hofree M, Choi D, Linton EL, Turkekul M, Bejnood A, et al. Regenerative potential of prostate luminal cells revealed by single-cell analysis. Science. 2020 May 1;368(6490):497–505. doi:10.1126/science.aay0267 PubMed PMID: 32355025; PubMed Central PMCID: PMC7313621.

66. He MX, Cuoco MS, Crowdis J, Bosma-Moody A, Zhang Z, Bi K, et al. Transcriptional mediators of treatment resistance in lethal prostate cancer. Nat Med. 2021 Mar;27(3):426–33. doi:10.1038/s41591-021-01244-6 PubMed PMID: 33664492; PubMed Central PMCID: PMC7960507.

67. Wang ZA, Toivanen R, Bergren SK, Chambon P, Shen MM. Luminal cells are favored as the cell of origin for prostate cancer. Cell Rep. 2014 Sep 11;8(5):1339–46. doi:10.1016/j.celrep.2014.08.002 PubMed PMID: 25176651; PubMed Central PMCID: PMC4163115.

68. Pitzen SP, Dehm SM. Basal epithelial cells in prostate development, tumorigenesis, and cancer progression. Cell Cycle. 2023;22(11):1303–18. doi:10.1080/15384101.2023.2206502 PubMed PMID: 37098827; PubMed Central PMCID: PMC10228417.

69. Tucci P, Agostini M, Grespi F, Markert EK, Terrinoni A, Vousden KH, et al. Loss of p63 and its microRNA-205 target results in enhanced cell migration and metastasis in prostate cancer. Proc Natl Acad Sci. 2012 Sep 18;109(38):15312–7. doi:10.1073/pnas.1110977109

70. Ekstrom TL, Rosok RM, Abdelrahman AM, Parassiadis C, Manjunath M, Dittrich MY, et al. Glucocorticoid receptor suppresses GATA6-mediated RNA polymerase II pause release to modulate classical subtype identity in pancreatic cancer. Gut. 2025 Jan 30;gutjnl-2024-334374. doi:10.1136/gutjnl-2024-334374 PubMed PMID: 39884837.

71. Dang TT, Esparza MA, Maine EA, Westcott JM, Pearson GW. ΔNp63α Promotes Breast Cancer Cell Motility through the Selective Activation of Components of the Epithelial-to-Mesenchymal Transition Program. Cancer Res. 2015 Sep 14;75(18):3925–35. doi:10.1158/0008-5472.CAN-14-3363

72. Lambert AW, Fiore C, Chutake Y, Verhaar ER, Strasser PC, Chen MW, et al. ΔNp63/p73 drive metastatic colonization by controlling a regenerative epithelial stem cell program in quasi-mesenchymal cancer stem cells. Dev Cell. 2022 Dec 19;57(24):2714–2730.e8. doi:10.1016/j.devcel.2022.11.015

73. Jin Z, Wang H, Tang R, Pan B, Lee HJ, Liu S, et al. GATA2 promotes castration-resistant prostate cancer development by suppressing IFN-β axis-mediated antitumor immunity. Oncogene. 2024 Aug;43(34):2595–610. doi:10.1038/s41388-024-03107-z

74. Gomes AM, Kurochkin I, Chang B, Daniel M, Law K, Satija N, et al. Cooperative Transcription Factor Induction Mediates Hemogenic Reprogramming. Cell Rep. 2018 Dec 4;25(10):2821–2835.e7. doi:10.1016/j.celrep.2018.11.032 PubMed PMID: 30517869; PubMed Central PMCID: PMC6571141.

75. Orlando KA, Grimm SA, Wade PA. Pioneering new enhancers by GATA3: role of facilitating transcription factors and chromatin remodeling. Nucleic Acids Res. 2025 Jun 6;53(11):gkaf473. doi:10.1093/nar/gkaf473 PubMed PMID: 40479714; PubMed Central PMCID: PMC12143596.

76. Paul S, Hagenbeek TJ, Tremblay J, Kameswaran V, Ong C, Liu C, et al. Cooperation between the Hippo and MAPK pathway activation drives acquired resistance to TEAD inhibition. Nat Commun. 2025 Feb 18;16:1743. doi:10.1038/s41467-025-56634-y PubMed PMID: 39966375; PubMed Central PMCID: PMC11836325.

77. Kaochar S, Rusin A, Foley C, Rajapakshe K, Robertson M, Skapura D, et al. Inhibition of GATA2 in prostate cancer by a clinically available small molecule. Endocr Relat Cancer. 2021 Oct 12;29(1):15–31. doi:10.1530/ERC-21-0085 PubMed PMID: 34636746; PubMed Central PMCID: PMC8634153.

