## Supplementary Figures for "Molecular switch mediates the glucocorticoid receptor transition from tumor suppressor to oncogene in the prostate"

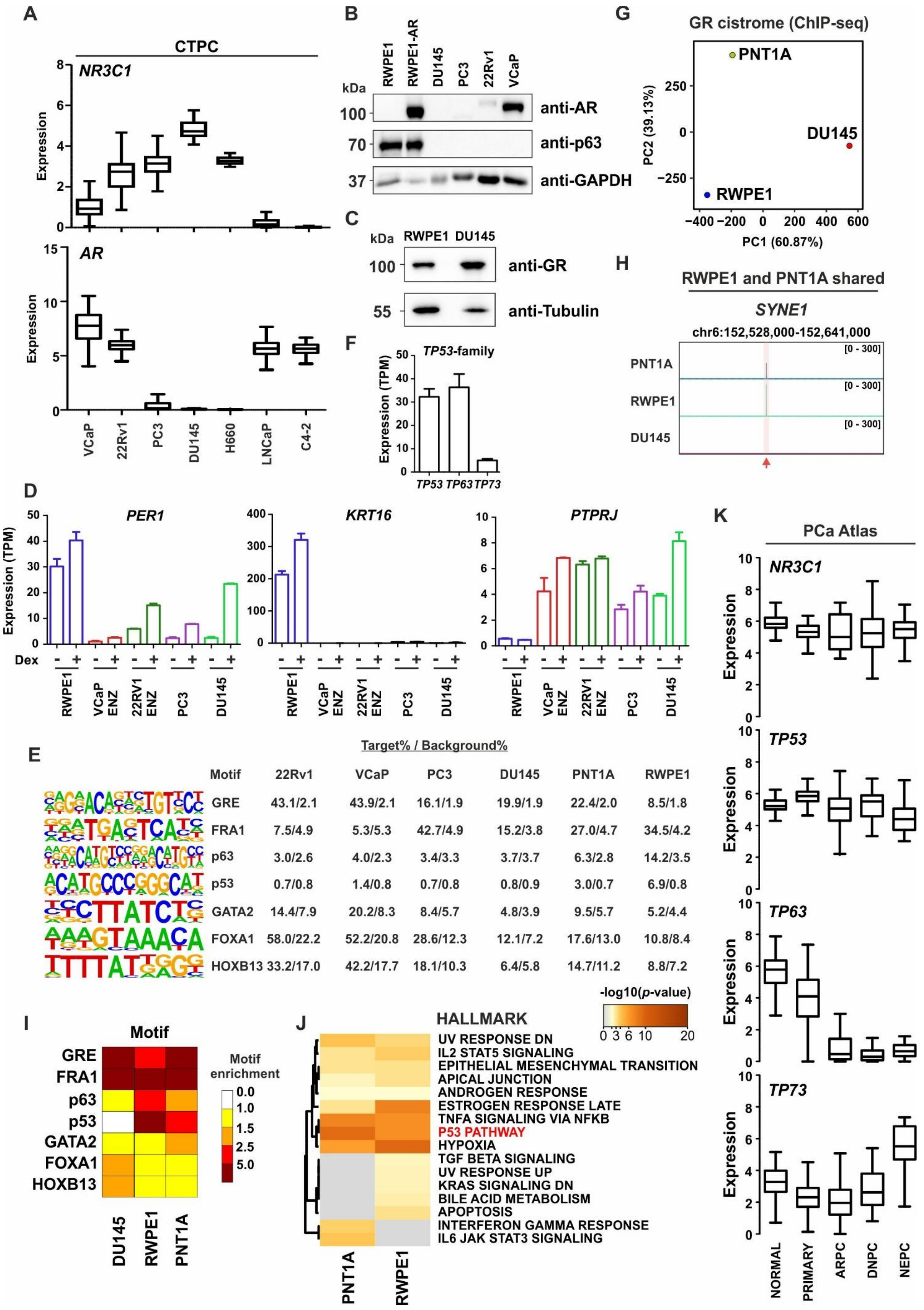
A
CTPC
NR3C1
Expression
AR
VCaP
22Rv1
PC3
DU145
H660
LNCaP
C4-2
B
kDa
RWPE1
RWPE1-AR
DU145
PC3
22Rv1
VCaP
100
anti-AR
70
anti-p63
37
anti-GAPDH
C
kDa
RWPE1 DU145
100
anti-GR
55
anti-Tubulin
F
TP53-family
Expression (TPM)
TP53 TP63 TP73
G
GR cistrome (ChIP-seq)
PNT1A
DU145
RWPE1
PC2 (39.13%)
PC1 (60.87%)
H
RWPE1 and PNT1A shared
SYNE1
chr6:152,528,000-152,641,000
PNT1A
RWPE1
DU145
[0 - 300]
D
PER1
KRT16
PTPRJ
Expression (TPM)
Dex
RWPE1
VCaP ENZ
22RV1 ENZ
PC3
DU145
E
Target% / Background%
Motif 22Rv1 VCaP PC3 DU145 PNT1A RWPE1
GRE 43.1/2.1 43.9/2.1 16.1/1.9 19.9/1.9 22.4/2.0 8.5/1.8
FRA1 7.5/4.9 5.3/5.3 42.7/4.9 15.2/3.8 27.0/4.7 34.5/4.2
p63 3.0/2.6 4.0/2.3 3.4/3.3 3.7/3.7 6.3/2.8 14.2/3.5
p53 0.7/0.8 1.4/0.8 0.7/0.8 0.8/0.9 3.0/0.7 6.9/0.8
GATA2 14.4/7.9 20.2/8.3 8.4/5.7 4.8/3.9 9.5/5.7 5.2/4.4
FOXA1 58.0/22.2 52.2/20.8 28.6/12.3 12.1/7.2 17.6/13.0 10.8/8.4
HOXB13 33.2/17.0 42.2/17.7 18.1/10.3 6.4/5.8 14.7/11.2 8.8/7.2
I
Motif
GRE
FRA1
p63
p53
GATA2
FOXA1
HOXB13
Motif enrichment
0.0
1.0
1.5
2.5
5.0
DU145
RWPE1
PNT1A
J
HALLMARK
-log10(p-value)
0 3 6 10 20
UV RESPONSE DN
IL2 STAT5 SIGNALING
EPITHELIAL MESENCHYMAL TRANSITION
APICAL JUNCTION
ANDROGEN RESPONSE
ESTROGEN RESPONSE LATE
TNFA SIGNALING VIA NFKB
P53 PATHWAY
HYPOXIA
TGF BETA SIGNALING
UV RESPONSE UP
KRAS SIGNALING DN
BILE ACID METABOLISM
APOPTOSIS
INTERFERON GAMMA RESPONSE
IL6 JAK STAT3 SIGNALING
PNT1A
RWPE1
K
PCa Atlas
NR3C1
TP53
TP63
TP73
Expression
NORMAL
PRIMARY
ARPC
DNPC
NEPC

**Supplementary Figure S1. Expression of *NR3C1* and *TP53*-family members in prostate cells.** (**A**) Expression of (upper) *NR3C1* and (lower) *AR* using Combined Transcriptome dataset of PCa Cell lines (CTPC) data. (**B**) Immunoblotting of AR, p63, and GAPDH protein levels in RWPE1, RWPE1-AR, DU145, PC3, 22Rv1, and VCaP cells. (**C**) Immunoblotting of GR and tubulin protein levels in RWPE1 and DU145 cells. (**D**) RNA-seq TPM values of *PER1*, *KRT16*, and *PTPRJ* expression from EtOH- and Dex-treated RWPE1, DU145, PC3, 22Rv1, and VCaP cells. 22Rv1 and VCaP cells had also been treated with Enzalutamide (ENZ). Bar graphs represent mean ±SD, n=3. (**E**) Known motif enrichment of specific motifs at GRBs in prostate and PCa cell lines. (**F**) RNA-seq TPM values of *TP53*, *TP63*, and *TP73* expression in RWPE1 cells. Bar graphs represent mean ±SD, n=3. (**G**) PCA plot of GR cistrome from RWPE1, PNT1A, and DU145 cells. (**H**) Genome browser track of GR ChIP-seq at *SYNE1* locus. Red arrow points specific peak at the locus. (**I**) Heatmap of specific motif enrichment at GRBs calculated as fold change over background. (**J**) Hallmark Gene Set pathway analysis of PNT1A and RWPE1 GR ChIP-seq data. Top 500 GRBs were selected for the analysis. Color scale represents $-\log_{10}$ p-value, and distinct pathways highlighted in red. (**K**) Expression of *NR3C1*, *TP53*, *TP63*, and *TP73* using PCa Atlas data from different subtypes of PCa.

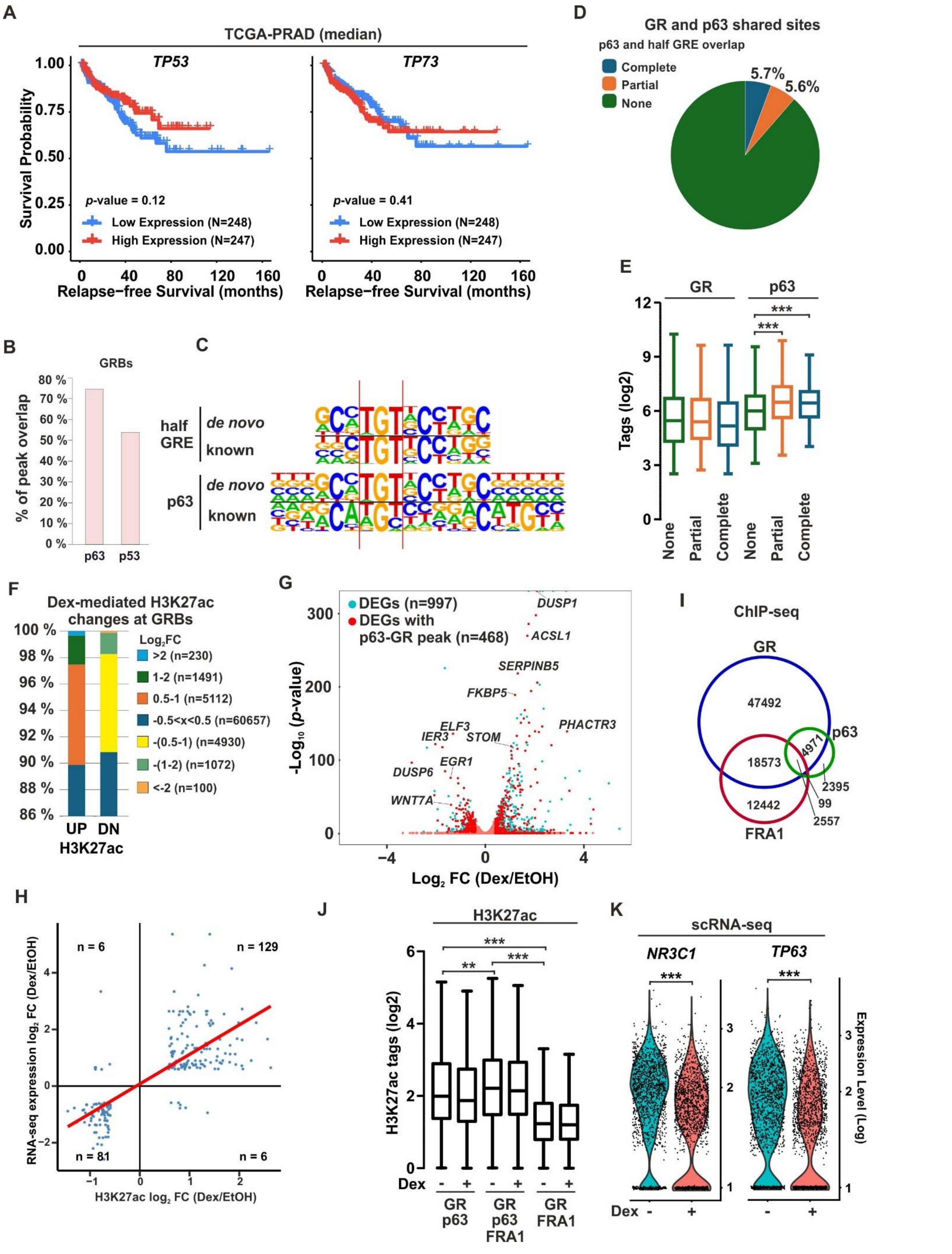

A
TCGA-PRAD (median)
TP53
TP73
Survival Probability
p-value = 0.12
p-value = 0.41
Low Expression (N=248)
High Expression (N=247)
Relapse−free Survival (months)
B
GRBs
% of peak overlap
p63
p53
C
half GRE
de novo
known
p63
D
GR and p63 shared sites
p63 and half GRE overlap
Complete
Partial
None
5.7%
5.6%
E
GR
p63
Tags (log2)
None
Partial
Complete
F
Dex-mediated H3K27ac changes at GRBs
Log2FC
>2 (n=230)
1-2 (n=1491)
0.5-1 (n=5112)
-0.5<x<0.5 (n=60657)
-(0.5-1) (n=4930)
-(1-2) (n=1072)
<-2 (n=100)
UP DN
H3K27ac
G
DEGs (n=997)
DEGs with p63-GR peak (n=468)
DUSP1
ACSL1
SERPINB5
FKBP5
PHACTR3
ELF3
IER3
STOM
EGR1
DUSP6
WNT7A
-Log10 (p-value)
Log2 FC (Dex/EtOH)
I
ChIP-seq
GR
p63
FRA1
47492
18573
4971
12442
2395
99
2557
H
RNA-seq expression log2 FC (Dex/EtOH)
H3K27ac log2 FC (Dex/EtOH)
n = 6
n = 129
n = 81
n = 6
J
H3K27ac
H3K27ac tags (log2)
Dex
GR p63
GR p63 FRA1
GR FRA1
K
scRNA-seq
NR3C1
TP63
Expression Level (Log)
Dex

**Supplementary Figure S2. GRBs overlap with p63 sites.** (**A**) Relapse-free survival of TCGA-PRAD patients expressing either high or low levels of (left) *TP53* or (right) *TP73*. PRAD, prostate adenocarcinoma; TCGA, the cancer genome atlas. (**B**) Overlap of p63 or p53 ChIP-seq peaks with GRBs in RWPE1 cells. (**C**) Shared nucleotides of the p63 and half GRE motifs using both known and de novo motifs on sites that contain both p63 and GRE motifs. Red lines highlight the conserved TGT in both motifs. (**D**) Percentage of p63-GR sites that have complete (blue), partial (orange), or no (green) p63 and half GRE motif overlap. (**E**) $Log_2$ tag enrichment of GR and p63 at the p63-GR sites with indicated overlap of p63 and half GRE motifs. (**F**) Dex-mediated H3K27ac changes at RWPE-1 GRBs. Bar graphs show the distribution of distinct $log_2$ fold changes ($log_2$ FC) of increased (UP) and decreased (DN) H3K27ac categories. (**G**) Volcano plot of Dex-mediated differential expressed genes (DEGs) in RWPE1 cells (n = 997, teal dots). Red dots indicate DEGs associated with GR-p63 binding sites (n = 468). (**H**) Relationship between Dex-mediated H3K27ac and gene expression changes at GR-p63-associated loci in RWPE1 cells. (**I**) Overlap of GR (blue), FRA1 (red), and p63 (green) ChIP-seq peaks in Dex-treated RWPE1 cells. (**J**) $Log_2$ tag enrichment of H3K27ac at the indicated sites with varying overlap with p63 and FRA1. (**K**) Violin plots of *NR3C1* and *TP63* expression in scRNA-seq data upon EtOH and Dex treatments (log1p). Statistical significance in box plots was calculated with One-way ANOVA with Bonferroni post hoc test, and in violin plots with Seurat. *, $p< 0.05$; **, $p< 0.01$; ***, $p< 0.001$.

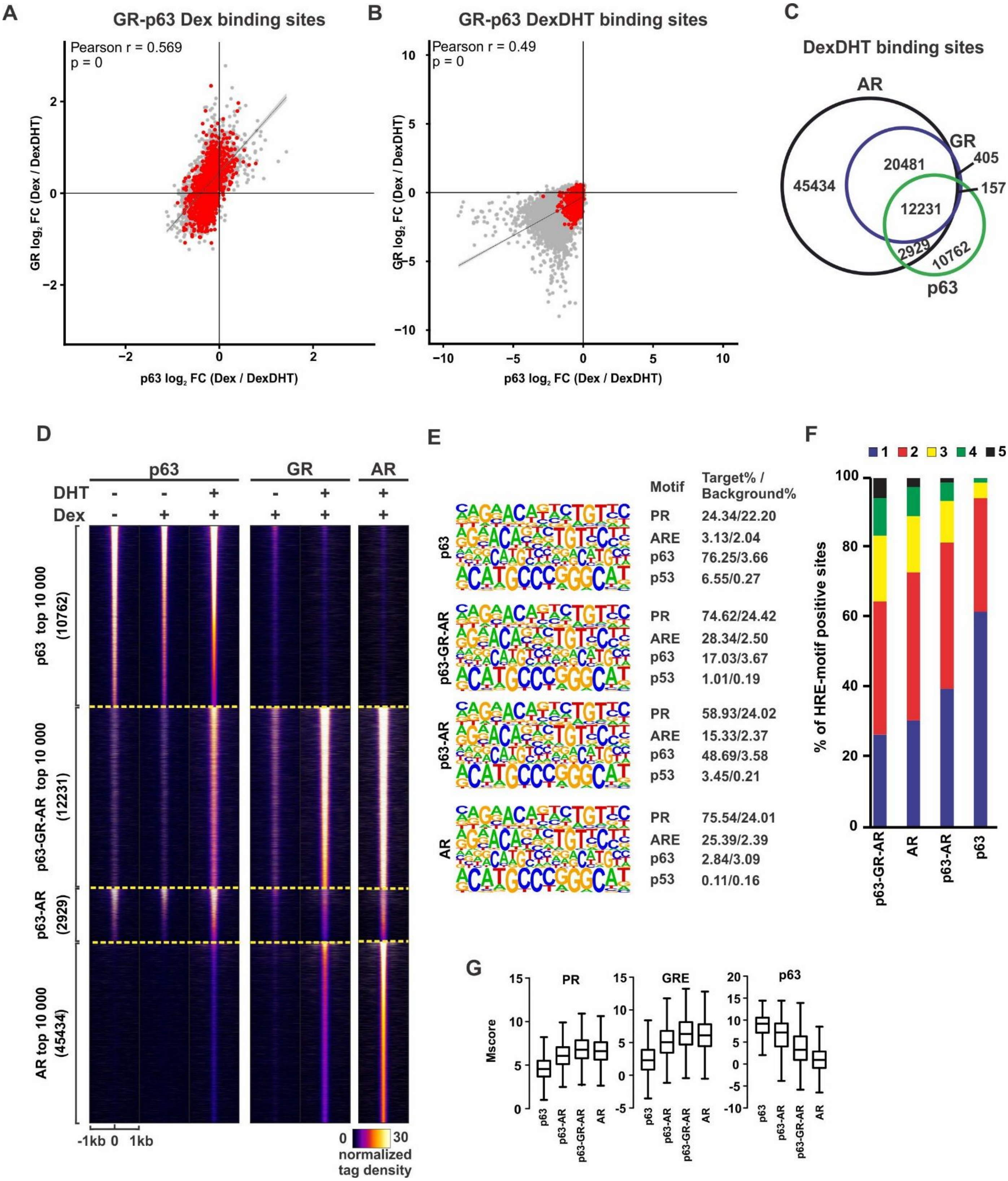

A
GR-p63 Dex binding sites
Pearson r = 0.569
p = 0
GR log2 FC (Dex / DexDHT)
p63 log2 FC (Dex / DexDHT)
B
GR-p63 DexDHT binding sites
Pearson r = 0.49
p = 0
C
DexDHT binding sites
AR
GR
p63
45434
20481
405
157
12231
2929
10762
D
p63
GR
AR
DHT
Dex
p63 top 10 000 (10762)
p63-GR-AR top 10 000 (12231)
p63-AR (2929)
AR top 10 000 (45434)
-1kb 0 1kb
normalized tag density
E
Motif
Target% / Background%
p63
PR 24.34/22.20
ARE 3.13/2.04
p63 76.25/3.66
p53 6.55/0.27
p63-GR-AR
PR 74.62/24.42
ARE 28.34/2.50
p63 17.03/3.67
p53 1.01/0.19
p63-AR
PR 58.93/24.02
ARE 15.33/2.37
p63 48.69/3.58
p53 3.45/0.21
AR
PR 75.54/24.01
ARE 25.39/2.39
p63 2.84/3.09
p53 0.11/0.16
F
% of HRE-motif positive sites
p63-GR-AR
AR
p63-AR
p63
G
Mscore
PR
GRE
p63

**Supplementary Figure S3. GR, p63 and AR binding sites overlap in RWPE1-AR cells.** (**A-B**) Correlation of GR and p63 $log_2$ fold change ($log_2$ FC) tag enrichment at GR-p63 binding sites in RWPE1-AR cells following Dex and DexDHT treatment (Dex / DexDHT). Sites represent p63-GR overlapping sites in (**A**) Dex- or (**B**) DexDHT-treated conditions. Red dots indicate GR-p63 peaks also present in RWPE1 cells. Pearson correlation coefficient and p-value are shown in the plot. (**C**) Overlap of GR (blue), AR (black), and p63 (green) ChIP-seq peaks in DexDHT-treated RWPE1-AR cells. (**D**) ChIP-seq profiles of p63, GR, and AR from RWPE1-AR cells at indicated overlapping binding site categories. Each heatmap represents ± 1 kb around the center of the peak, and binding intensity (tags per bp per site) is noted below on a linear scale. (**E**) Known motif enrichment of specific motifs at indicated binding site categories. (**F**) The number of HRE-motif (PR motif) positive at indicated binding site categories. (**G**) Box plot of PR, GRE, and p63 motif scores at indicated binding site categories.

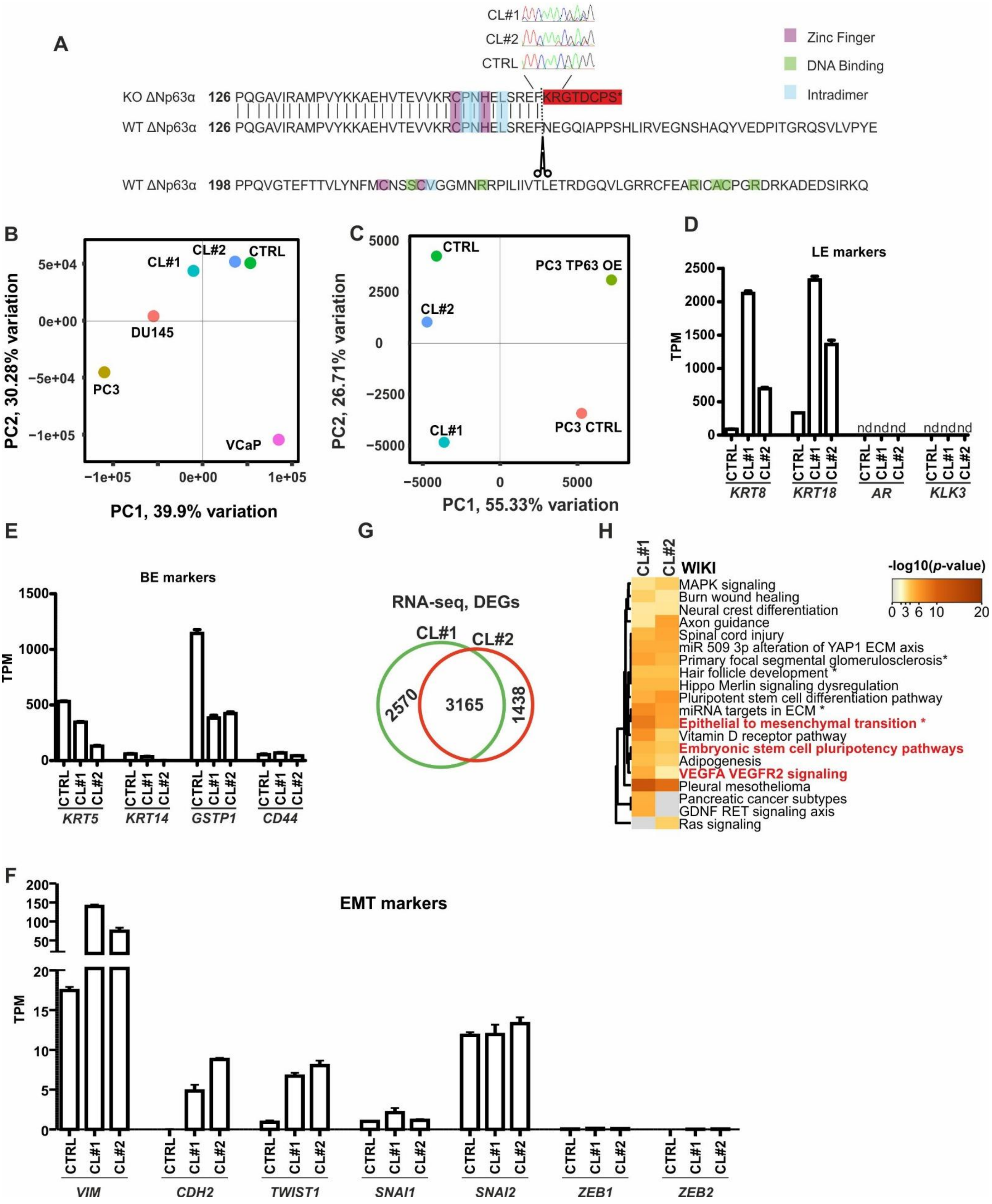
A
CL#1
CL#2
CTRL
Zinc Finger
DNA Binding
Intradimer
KO ΔNp63α 126 PQGAVIRAMPVYKKAEHVTEVVKRCPNHELSREFKRGTDCPS*
WT ΔNp63α 126 PQGAVIRAMPVYKKAEHVTEVVKRCPNHELSREFNEGQIAPPSHLIRVEGNSHAQYVEDPITGRQSVLVPYE
WT ΔNp63α 198 PPQVGTEFTTVLYNFMCNSSCVGGMNRRPILIIVTLETRDGQVLGRRCFEARICACPGRDRKADEDSIRKQ
B
PC1, 39.9% variation
PC2, 30.28% variation
C
PC1, 55.33% variation
PC2, 26.71% variation
PC3 TP63 OE
PC3 CTRL
D
LE markers
TPM
KRT8
KRT18
AR
KLK3
E
BE markers
KRT5
KRT14
GSTP1
CD44
G
RNA-seq, DEGs
2570
3165
1438
H
WIKI
-log10(p-value)
MAPK signaling
Burn wound healing
Neural crest differentiation
Axon guidance
Spinal cord injury
miR 509 3p alteration of YAP1 ECM axis
Primary focal segmental glomerulosclerosis*
Hair follicle development *
Hippo Merlin signaling dysregulation
Pluripotent stem cell differentiation pathway
miRNA targets in ECM *
Epithelial to mesenchymal transition *
Vitamin D receptor pathway
Embryonic stem cell pluripotency pathways
Adipogenesis
VEGFA VEGFR2 signaling
Pleural mesothelioma
Pancreatic cancer subtypes
GDNF RET signaling axis
Ras signaling
F
EMT markers
VIM
CDH2
TWIST1
SNAI1
SNAI2
ZEB1
ZEB2

**Supplementary Figure S4. CRISPR-Cas9-mediated depletion of p63.** (**A**) CRISPR-Cas9 induced frame shift in p63 amino acid sequence leads to an early stop codon in p63KO cells. Crucial amino acids for Zinc finger (purple), DNA binding (green), and intradimer (blue) shown in the sequence. (**B**) PCA plot of DU145, PC3, VCaP, RWPE1 CTRL and -p63KO cell line transcriptomes. (**C**) PCA plot of RWPE1 CTRL and -p63KO, and PC3 CTRL and -p63OE cell line transcriptomes. (**D**) RNA-seq TPM values of LE marker genes *KRT8*, *KRT18*, *AR*, and *KLK3* from CTRL and p63KO cells. (**E**) RNA-seq TPM values of BE marker genes *KRT5*, *KRT14*, *GSTP1*, and *CD44* from CTRL and p63KO cells. (**F**) RNA-seq TPM values of EMT marker genes *VIM*, *CDH2*, *TWIST1*, *SNAI1*, *SNAI2*, *ZEB1*, and *ZEB2* from CTRL and p63KO cells (**G**) Overlap of DEGs in CL#1 (green) and CL#2 (red) p63KO cells. (**H**) WIKI pathway analysis of CL#1 and CL#2 DEGs with specific pathways highlighted in red.

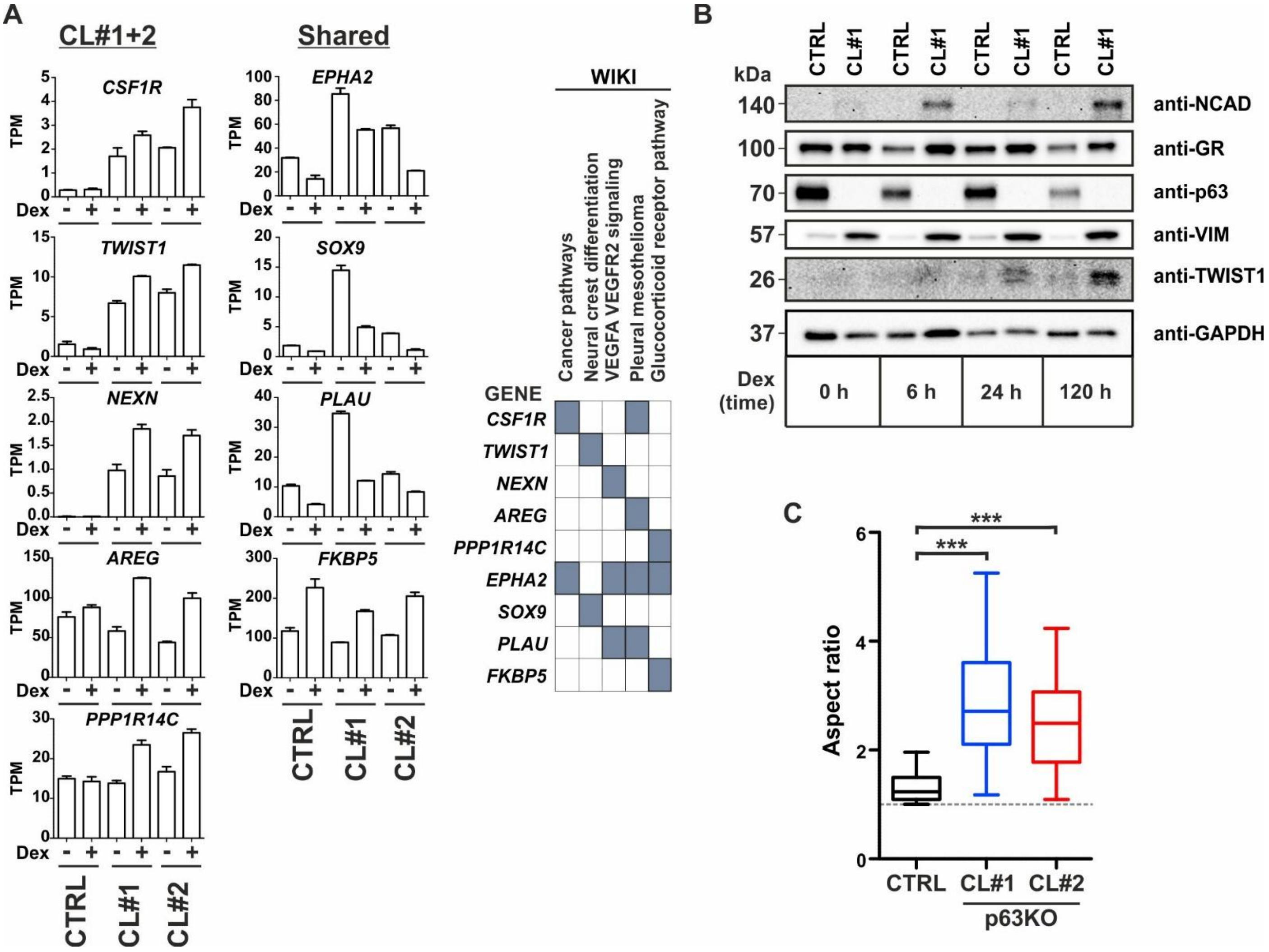


**Supplementary Figure S5. p63KO cells display increased EMT markers.** (**A**) (left) RNA-seq TPM values of specific Dex-regulated transcripts from CTRL and p63KO cells in CL#1+2 and Shared categories. (right) WIKI pathway association of indicated Dex-regulated genes. Bar graphs represent mean ±SD, n=2. (**B**) Immunoblot of N-cadherin (NCAD), GR, p63, Vimentin (VIM), TWIST1, and GAPDH in CTRL and p63KO (CL#1) cells following dexamethasone (Dex) treatment for 0 h, 6 h, 24 h (1 day), and 120 h (5 day). (**C**) Aspect ratio from CTRL and p63KO cells, displaying the ratio of cell length to width (n=37-41). Statistical significance in box plots was calculated with One-way ANOVA with Bonferroni post hoc test. *, $p < 0.05$; **, $p < 0.01$; ***, $p < 0.001$.

A

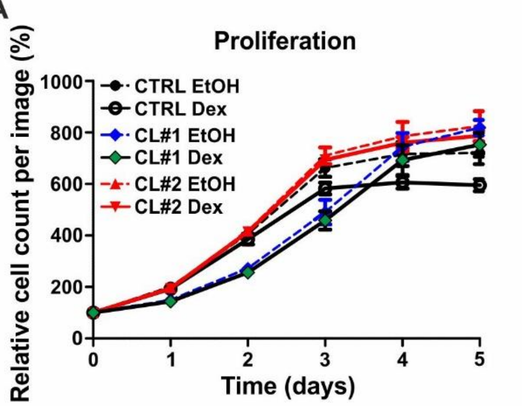


B

**Proliferation**

| CTRL EtOH vs CTRL Dex | | | | |
|---|---|---|---|---|
| Time (days) | Difference | t | P value | Summary |
| 3 | -79.36 | 4.387 | P<0.001 | *** |
| 4 | -109.8 | 6.068 | P<0.001 | *** |
| 5 | -125.1 | 6.913 | P<0.001 | *** |

| CL#1 EtOH vs CL#1 Dex | | | | |
|---|---|---|---|---|
| Time (days) | Difference | t | P value | Summary |
| 3 | -32.32 | 1.684 | P > 0.05 | ns |
| 4 | -50.4 | 2.626 | P > 0.05 | ns |
| 5 | -65.95 | 3.437 | P<0.01 | ** |

| CL#2 EtOH vs CL#2 Dex | | | | |
|---|---|---|---|---|
| Time (days) | Difference | t | P value | Summary |
| 3 | -16.13 | 0.8918 | P > 0.05 | ns |
| 4 | -23.84 | 1.318 | P > 0.05 | ns |
| 5 | -37.34 | 2.064 | P > 0.05 | ns |

C

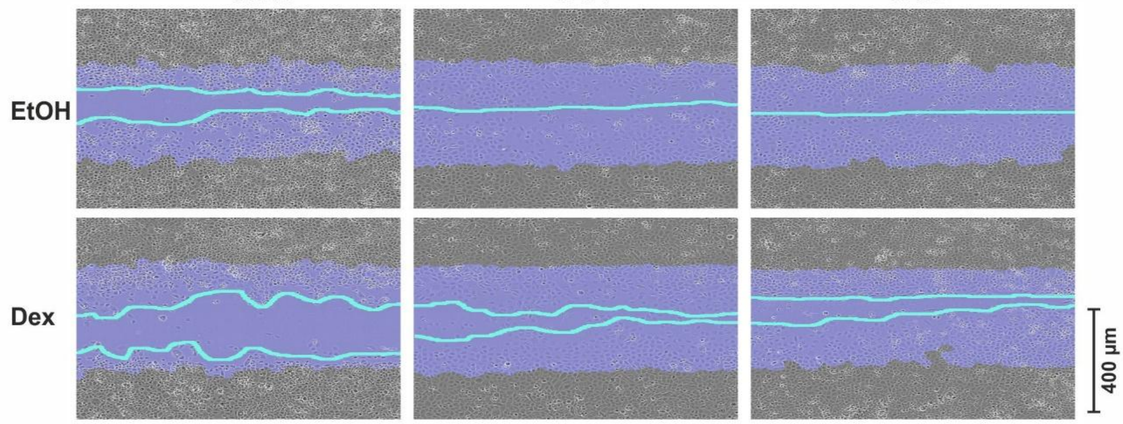


D

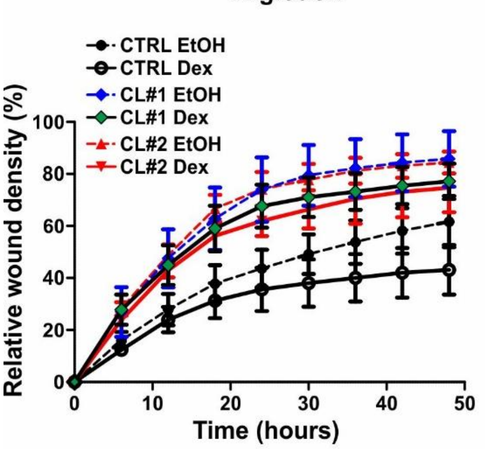


E

**Migration**

| CTRL EtOH vs CTRL Dex | | | | |
|---|---|---|---|---|
| Time (h) | Difference | t | P value | Summary |
| 36 | -13.76 | 2.784 | P > 0.05 | ns |
| 42 | -16.12 | 3.261 | P < 0.05 | * |
| 48 | -18.49 | 3.74 | P<0.01 | ** |

| CL#1 EtOH vs CL#1 Dex | | | | |
|---|---|---|---|---|
| Time (h) | Difference | t | P value | Summary |
| 36 | -8.996 | 1.82 | P > 0.05 | ns |
| 42 | -8.815 | 1.783 | P > 0.05 | ns |
| 48 | -8.602 | 1.74 | P > 0.05 | ns |

| CL#2 EtOH vs CL#2 Dex | | | | |
|---|---|---|---|---|
| Time (h) | Difference | t | P value | Summary |
| 36 | -10.78 | 2.18 | P > 0.05 | ns |
| 42 | -10.01 | 2.025 | P > 0.05 | ns |
| 48 | -9.733 | 1.969 | P > 0.05 | ns |

G

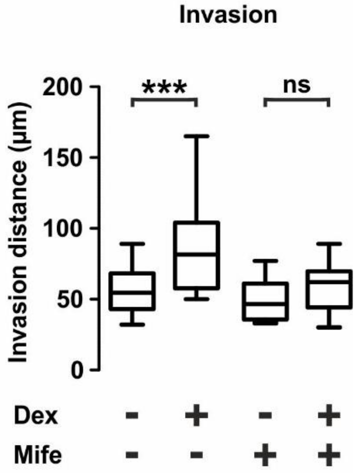


F

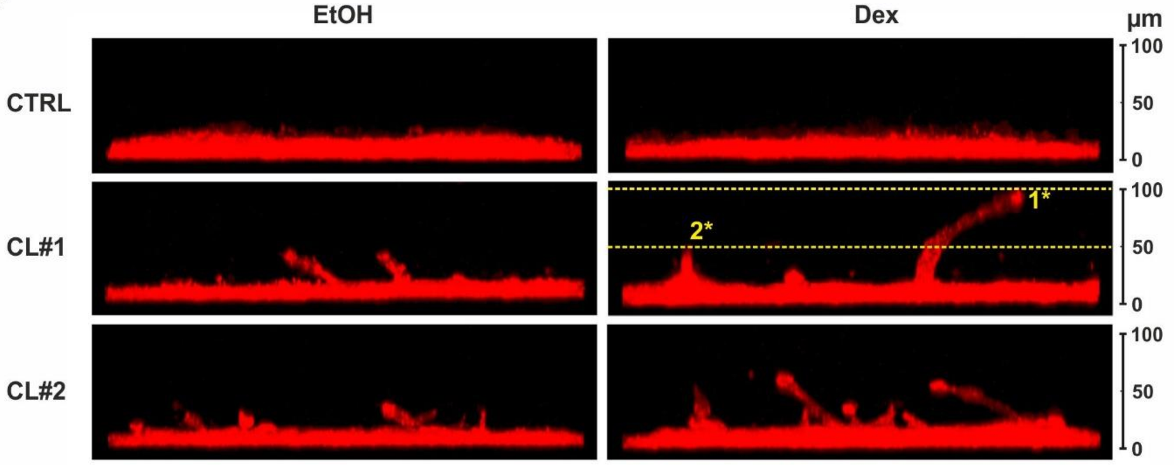

**Supplementary Figure S6. Analysis of proliferation, migration and invasion of p63KO cells.** (**A**) Relative cell proliferation in EtOH (dashed line) and Dex (solid line) treated CTRL (black), p63KO (CL#1 EtOH, blue; CL#1 Dex, green; CL#2 EtOH and Dex, red) cells. Data is normalized to day 0 and represents the mean ±SD of 10 images per day. (**B**) Summary table of the statistical comparison between EtOH- and Dex-treated CTRL and p63KO cell proliferation at the three last timepoints. (**C**) Examples of wound healing assay, with scale bar, 400 µm. Blue area depicts the initial wound, with the teal line depicting the confluence of the cells after 48 h. (**D**) Relative cell migration in EtOH (dashed line) and Dex (solid line) treated CTRL (black), p63KO (CL#1 EtOH, blue; CL#1 Dex, green; CL#2 EtOH and Dex, red) cells. Data is normalized to day 0 and represents the mean ±SD of 8 images per day. (**E**) Summary table of the statistical comparison between EtOH- and Dex-treated CTRL and p63KO cell migration at the three last timepoints. (**F**) Examples of 3D gel invasion, with scale bar, 100 µm. Two of the highest invasion points per cell were utilized (n=36-52). Yellow dashed lines in CL#1 Dex-treated cells depict an example of the highest invasion points. (**G**) 3D gel invasion of CTRL and p63KO cells after day 6 of EtOH, Dex, or mifepristone (Mife) treatments. Two of the highest invasion points were utilized (n=22). Statistical significance calculated with Two-way ANOVA with Bonferroni post hoc test. *, $p < 0.05$; **, $p < 0.01$; ***, $p < 0.001$; ns, non-significant.

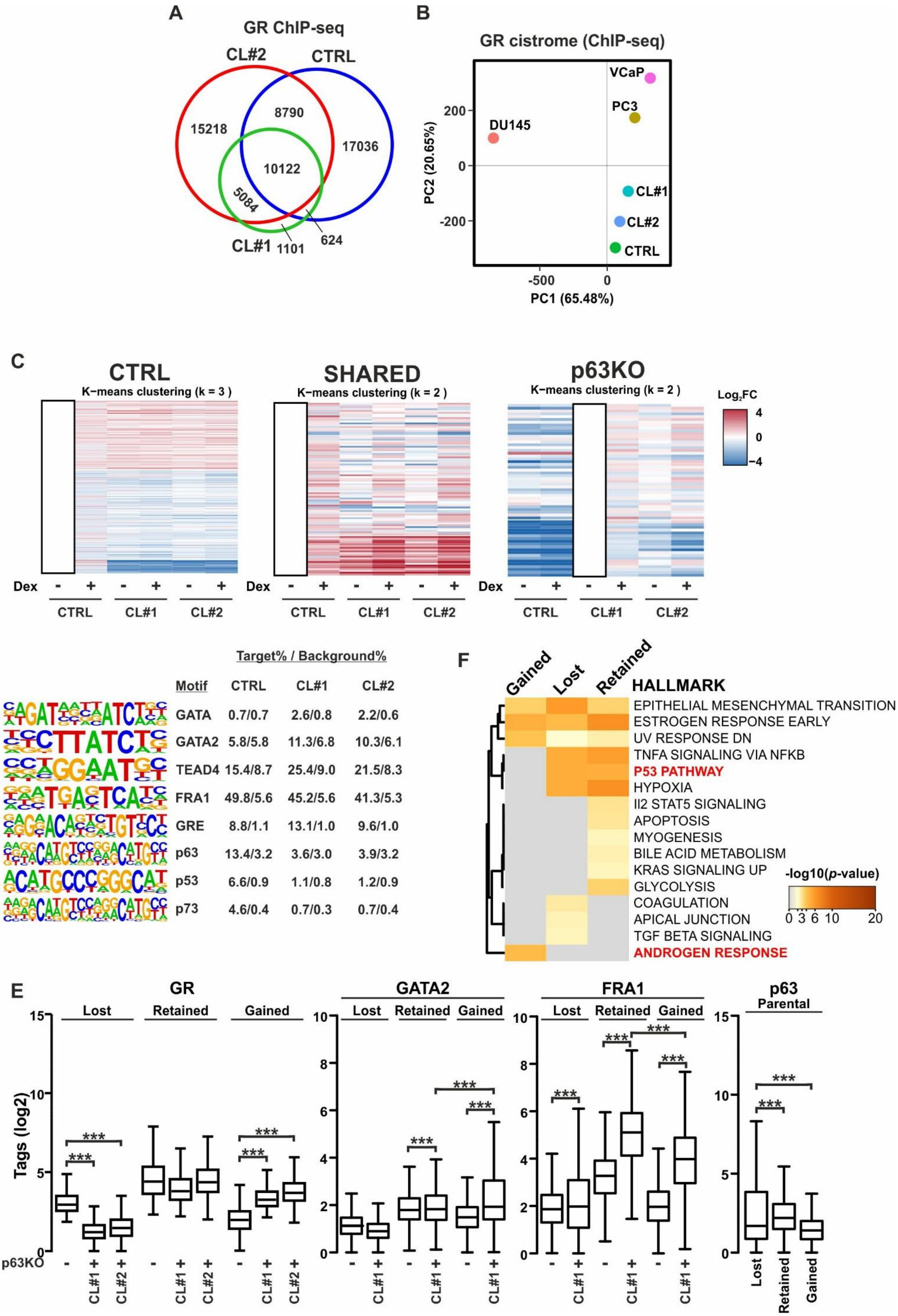
A
GR ChIP-seq
CL#2
CTRL
CL#1
15218
8790
17036
10122
5084
1101
624
B
GR cistrome (ChIP-seq)
VCaP
PC3
DU145
CL#1
CL#2
CTRL
PC2 (20.65%)
PC1 (65.48%)
C
CTRL
K−means clustering (k = 3 )
SHARED
K−means clustering (k = 2 )
p63KO
K−means clustering (k = 2 )
Log2FC
Dex
CTRL
CL#1
CL#2
D
Target% / Background%
Motif CTRL CL#1 CL#2
GATA 0.7/0.7 2.6/0.8 2.2/0.6
GATA2 5.8/5.8 11.3/6.8 10.3/6.1
TEAD4 15.4/8.7 25.4/9.0 21.5/8.3
FRA1 49.8/5.6 45.2/5.6 41.3/5.3
GRE 8.8/1.1 13.1/1.0 9.6/1.0
p63 13.4/3.2 3.6/3.0 3.9/3.2
p53 6.6/0.9 1.1/0.8 1.2/0.9
p73 4.6/0.4 0.7/0.3 0.7/0.4
F
Gained
Lost
Retained
HALLMARK
EPITHELIAL MESENCHYMAL TRANSITION
ESTROGEN RESPONSE EARLY
UV RESPONSE DN
TNFA SIGNALING VIA NFKB
P53 PATHWAY
HYPOXIA
Il2 STAT5 SIGNALING
APOPTOSIS
MYOGENESIS
BILE ACID METABOLISM
KRAS SIGNALING UP
GLYCOLYSIS
COAGULATION
APICAL JUNCTION
TGF BETA SIGNALING
ANDROGEN RESPONSE
-log10(p-value)
E
GR
GATA2
FRA1
p63
Parental
Lost
Retained
Gained
Tags (log2)
p63KO
CL#1
CL#2
***

**Supplementary Figure S7. GR binding and interactions are reorganized upon p63KO.** (**A**) Overlap of GR ChIP-seq peaks in Dex-treated CTRL (blue), p63KO (CL#1, green; CL#2, red) cells. (**B**) PCA plot of DU145, PC3, VCaP, RWPE1 CTRL and -p63KO cell line GR cistromes. (**C**) Heatmaps showing $\log_2$ fold change ($\log_2$ FC) expression changes in GR-regulated genes following Dex-treatment. Dex-regulated genes in Control, p63KO, and in both were clustered by k-means clustering. White box indicates the reference treatment for each heatmap. (**D**) Known motif enrichment of specific motifs at GRBs in CTRL and p63KO cells. (**E**) $\text{Log}_2$ tag enrichment of GR, GATA2, and FRA1 in p63-intact (p63KO -) and p63KO (p63KO +) cells, and p63 in parental RWPE1 cells at indicated p63KO GRBs. (**F**) Hallmark Gene Sets pathway analysis of p63KO GRBs. Top 500 GRBs were selected for the analysis. Color scale represents $-\log_{10}$ p-value, and distinct pathways highlighted in red. Statistical significance in box plots was calculated with One-way ANOVA with Bonferroni post hoc test. *, $p < 0.05$; **, $p < 0.01$; ***, $p < 0.001$.

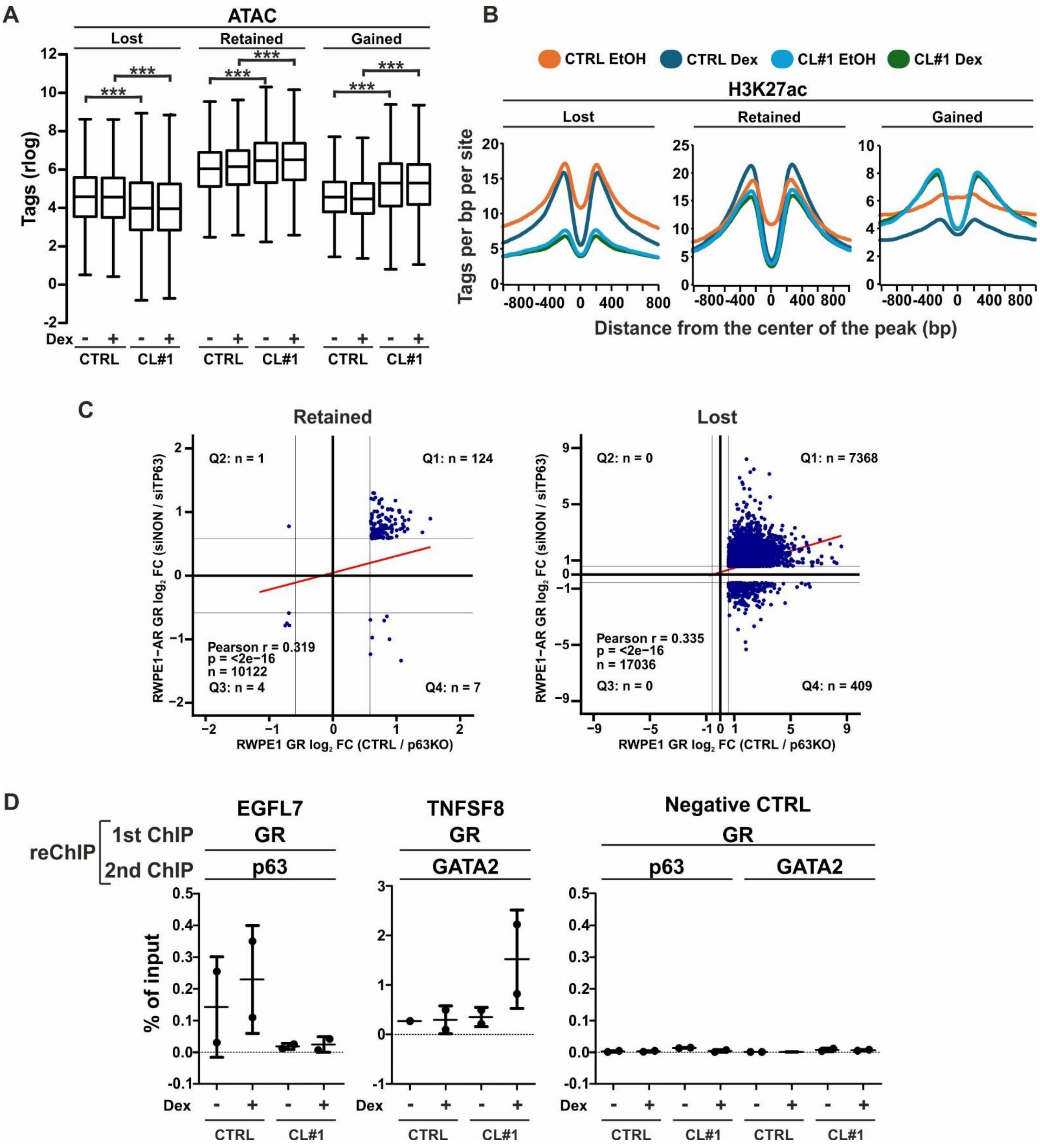
A
ATAC
Lost
Retained
Gained
Tags (rlog)
Dex
CTRL
CL#1
B
CTRL EtOH
CTRL Dex
CL#1 EtOH
CL#1 Dex
H3K27ac
Lost
Retained
Gained
Tags per bp per site
Distance from the center of the peak (bp)
C
Retained
Q2: n = 1
Q1: n = 124
Pearson r = 0.319
p = <2e−16
n = 10122
Q3: n = 4
Q4: n = 7
RWPE1−AR GR log2 FC (siNON / siTP63)
RWPE1 GR log2 FC (CTRL / p63KO)
Lost
Q2: n = 0
Q1: n = 7368
Pearson r = 0.335
p = <2e−16
n = 17036
Q3: n = 0
Q4: n = 409
D
reChIP
1st ChIP
2nd ChIP
EGFL7
GR
p63
TNFSF8
GR
GATA2
Negative CTRL
GR
p63
GATA2
% of input
Dex
CTRL
CL#1

**Supplementary Figure S8. GR binding and interactions are reorganized upon p63KO.** (**A**) Variance stabilized and transformed (rlog) tag enrichment of ATAC-seq in CTRL and p63KO cells at indicated p63KO GRBs. (**B**) H3K27ac rlog tag enrichment in CTRL and p63KO cells at indicated p63KO GRBs. Histograms represent ± 800 bp around the center of the peak, and enrichment intensity represents tags per bp per site. (**C**) Relationship between GR $log_2$ fold change ($log_2$ FC) tag enrichment in Dex-treated cells at (left) retained and (right) lost GRBs in CTRL versus p63KO cells (x-axis; CTRL / p63KO) and in RWPE1-AR cells treated with control siRNA (siNON) or p63 siRNA (siTP63) (y-axis; siNON / siTP63). Vertical and horizontal lines indicate a $-1.5 > FC < 1.5$ threshold. (**D**) Sequential (re)ChIP-qPCR analysis of GR+p63 binding on *EGFL7* loci, GR+GATA2 binding on *TNFSF8* loci, and binding of both negative control loci in EtOH or Dex treated CTRL or p63KO cells. Line and whiskers represent average ± SD, n=2. Statistical significance in box plots was calculated with One-way ANOVA with Bonferroni post hoc test. *, $p < 0.05$; **, $p < 0.01$; ***, $p < 0.001$.

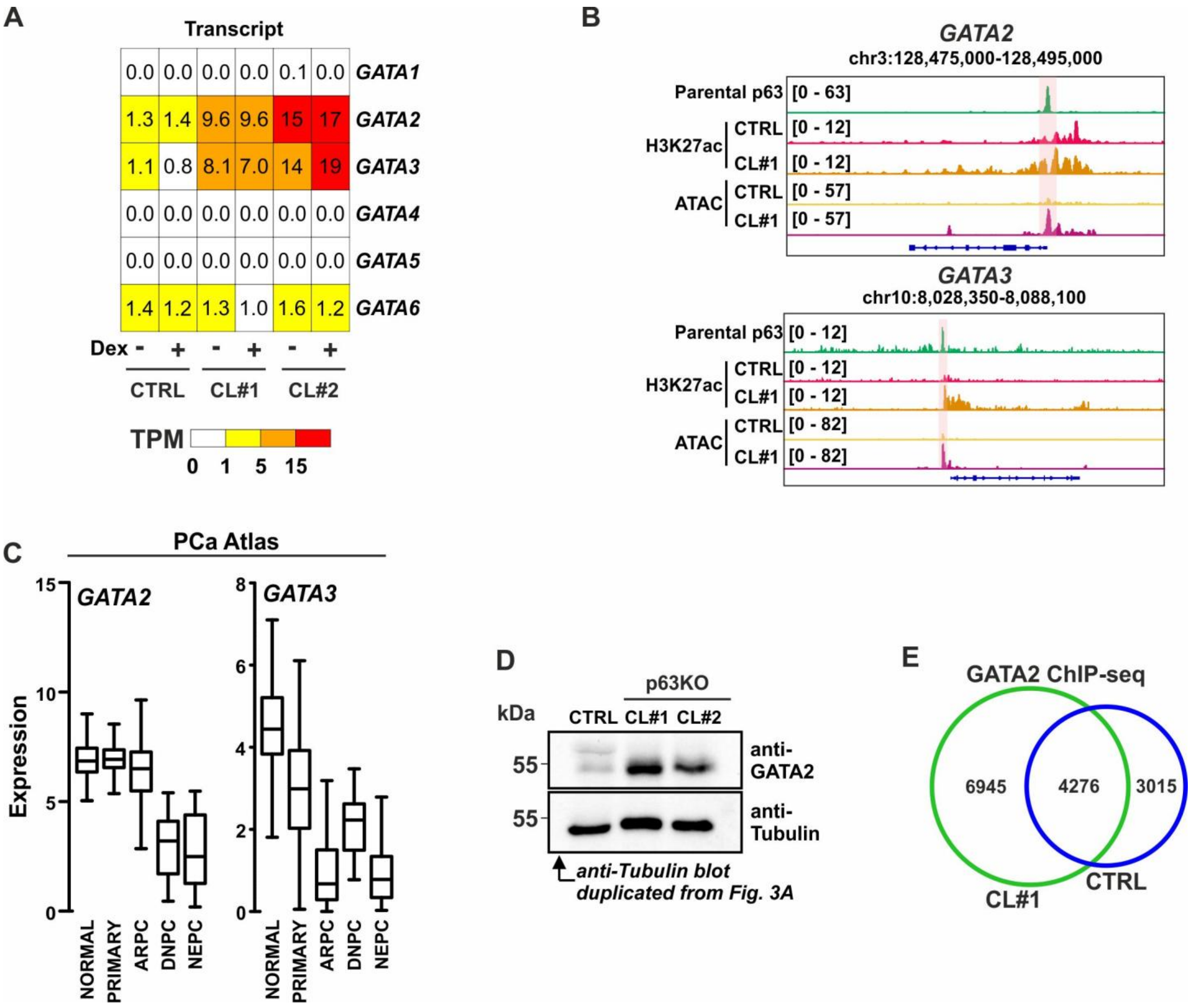


**Supplementary Figure S9. GATA2 expression and GR interactions are enhanced in p63KO cells.** (**A**) Heatmap of *GATA*-family member TPM values in CTRL and p63KO cells. (**B**) Genome browser tracks of p63 ChIP-seq from parental RWPE1 cells, and H3K27ac ChIP-seq and ATAC-seq from CTRL and p63KO cells at GATA2 and GATA3 loci. (**C**) Expression of *GATA2* and *GATA3* using PCa Atlas data from different subtypes of PCa. (**D**) Immunoblotting of GATA2 and tubulin protein levels in CTRL and p63KO cells. Due to the exact same protein samples, the tubulin immunoblot is copy from Figure 3A. (**E**) Overlap of GATA2 ChIP-seq peaks in CTRL (blue) and p63KO (green) cells.

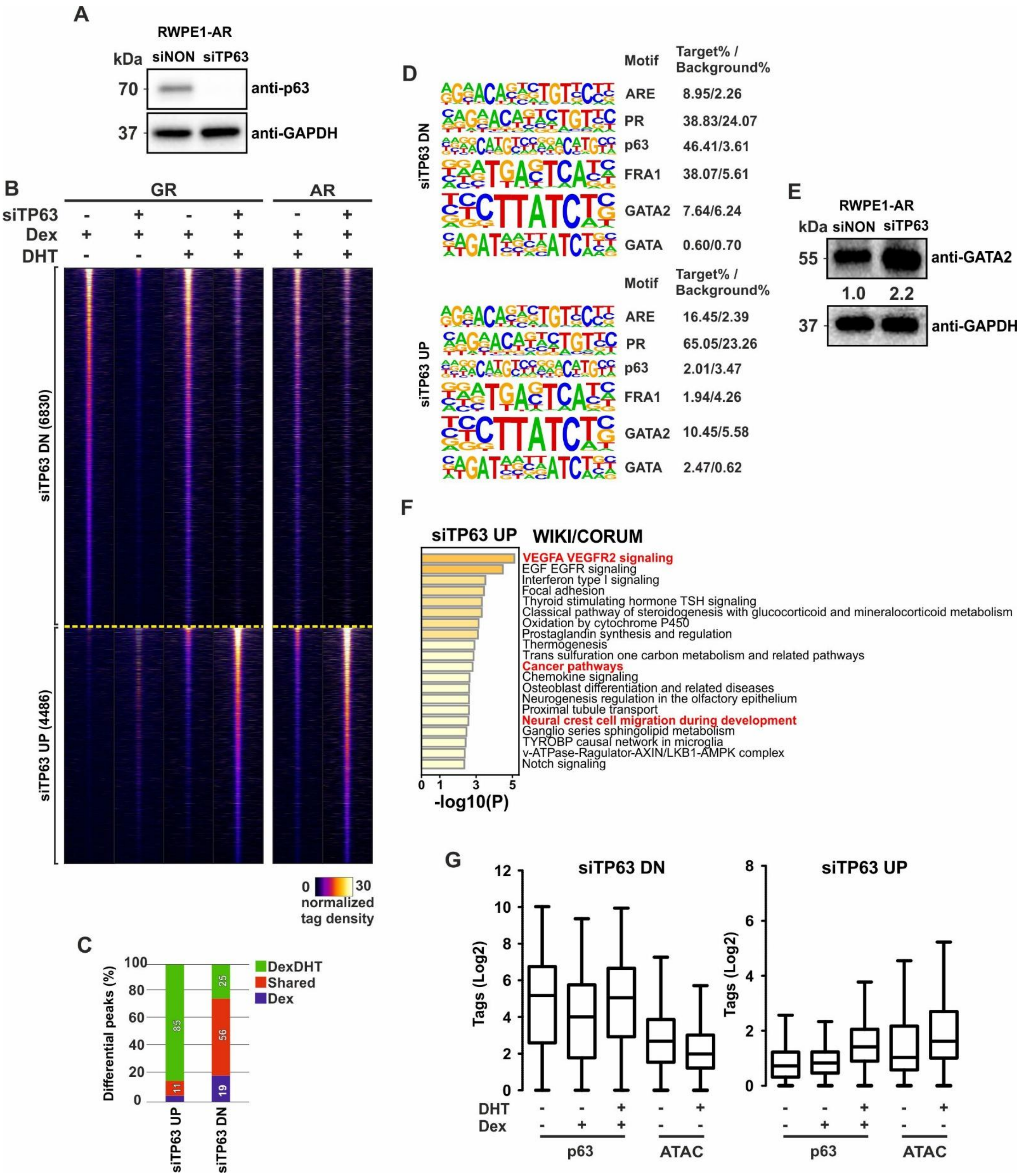

A
RWPE1-AR
kDa siNON siTP63
70 anti-p63
37 anti-GAPDH
B
GR
AR
siTP63 - + - + - +
Dex + + + + + +
DHT - - + + + +
siTP63 DN (6830)
siTP63 UP (4486)
0 30
normalized tag density
C
Differential peaks (%)
DexDHT
Shared
Dex
85
11
25
56
19
siTP63 UP
siTP63 DN
D
Motif Target% / Background%
siTP63 DN
ARE 8.95/2.26
PR 38.83/24.07
p63 46.41/3.61
FRA1 38.07/5.61
GATA2 7.64/6.24
GATA 0.60/0.70
siTP63 UP
ARE 16.45/2.39
PR 65.05/23.26
p63 2.01/3.47
FRA1 1.94/4.26
GATA2 10.45/5.58
GATA 2.47/0.62
E
RWPE1-AR
kDa siNON siTP63
55 anti-GATA2
1.0 2.2
37 anti-GAPDH
F
siTP63 UP WIKI/CORUM
VEGFA VEGFR2 signaling
EGF EGFR signaling
Interferon type I signaling
Focal adhesion
Thyroid stimulating hormone TSH signaling
Classical pathway of steroidogenesis with glucocorticoid and mineralocorticoid metabolism
Oxidation by cytochrome P450
Prostaglandin synthesis and regulation
Thermogenesis
Trans sulfuration one carbon metabolism and related pathways
Cancer pathways
Chemokine signaling
Osteoblast differentiation and related diseases
Neurogenesis regulation in the olfactory epithelium
Proximal tubule transport
Neural crest cell migration during development
Ganglio series sphingolipid metabolism
TYROBP causal network in microglia
v-ATPase-Ragulator-AXIN/LKB1-AMPK complex
Notch signaling
-log10(P)
G
siTP63 DN
siTP63 UP
Tags (Log2)
DHT - - + - +
Dex - + + - -
p63
ATAC

**Supplementary Figure S10. RWPE1-AR GR cistrome is remodeled by p63 depletion.** (**A**) Immunoblot of p63 and GAPDH protein levels in control siRNA (siNON) or p63 siRNA (siTP63)-treated RWPE1-AR cells. (**B**) ChIP-seq profiles of GR and AR from RWPE1-AR cells at GRBs that are decreased (siTP63 DN) or increased (siTP63 UP) following siTP63 treatment. Each heatmap represents ± 1 kb around the center of the peak, and binding intensity (tags per bp per site) is noted below on a linear scale. (**C**) Stacked bar graph show the percentage proportion of differential siTP63 UP and DN GRBs at Dex-, DexDHT-, and shared binding site categories. Differential enrichment defined as a ≥2-fold change in GR tag counts. (**D**) Known motif enrichment of specific motifs at siTP63 UP and DN GRBs in RWPE1-AR cells. (**E**) Immunoblot of GATA2 and GAPDH protein levels in siNON- and siTP63-treated RWPE1-AR cells. Quantification of the GATA2 displayed below the immunoblot with each normalized to GAPDH and compared to GATA2 siNON sample. (**F**) WIKI and CORUM pathway analysis of siTP63 UP sites. Top 500 GRBs were selected for the analysis. Distinct pathways highlighted in red. (**G**) p63 ChIP-seq and ATAC-seq tag enrichment on siTP63 UP and DN GRBs in RWPE1-AR cells treated with EtOH, Dex and/or DHT.

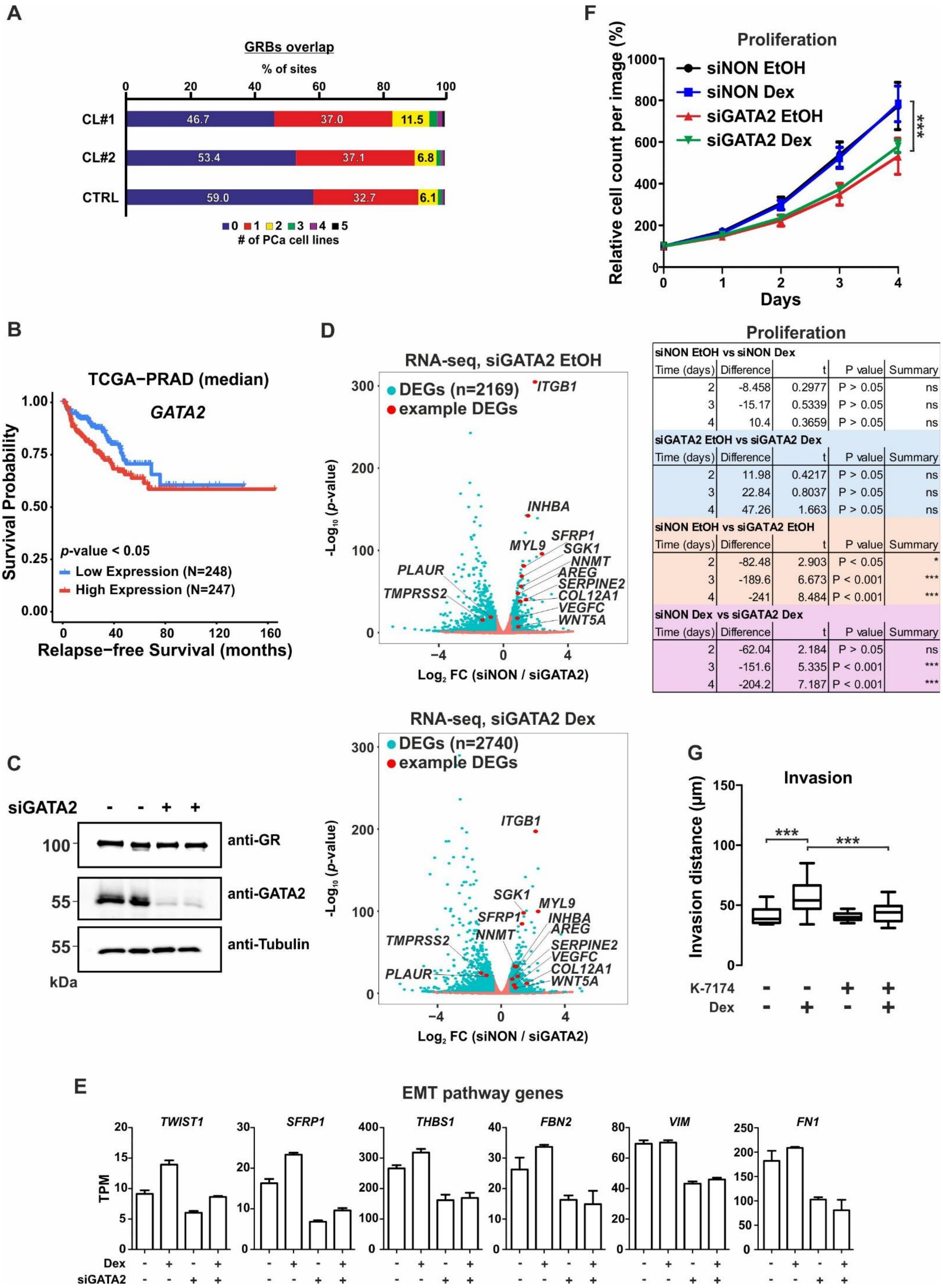


Proliferation

| Time (days) | Difference | t | P value | Summary |
|---|---|---|---|---|
| **siNON EtOH vs siNON Dex** | | | | |
| 2 | -8.458 | 0.2977 | P > 0.05 | ns |
| 3 | -15.17 | 0.5339 | P > 0.05 | ns |
| 4 | 10.4 | 0.3659 | P > 0.05 | ns |
| **siGATA2 EtOH vs siGATA2 Dex** | | | | |
| 2 | 11.98 | 0.4217 | P > 0.05 | ns |
| 3 | 22.84 | 0.8037 | P > 0.05 | ns |
| 4 | 47.26 | 1.663 | P > 0.05 | ns |
| **siNON EtOH vs siGATA2 EtOH** | | | | |
| 2 | -82.48 | 2.903 | P < 0.05 | * |
| 3 | -189.6 | 6.673 | P < 0.001 | *** |
| 4 | -241 | 8.484 | P < 0.001 | *** |
| **siNON Dex vs siGATA2 Dex** | | | | |
| 2 | -62.04 | 2.184 | P > 0.05 | ns |
| 3 | -151.6 | 5.335 | P < 0.001 | *** |
| 4 | -204.2 | 7.187 | P < 0.001 | *** |

**Supplementary Figure S11. GATA2-dependent activities in p63KO cells.** (**A**) GR ChIP-seq overlap of CTRL and p63KO cells with five PCa cell line GR cistrome (VCaP, 22Rv1, PC3, DU145, LREX). (**B**) Relapse-free survival of TCGA-PRAD patients expressing either high or low levels of *GATA2*. (**C**) Immunoblotting of GR, GATA2 and tubulin protein levels in siNON- and siGATA2-treated p63KO cells. (**D**) Volcano plot of siGATA2 DEGs (blue) in EtOH (upper) and Dex (lower) treated p63KO cells. Data depicts $\log_2$ fold change values of siNON/siGATA2 (x-axis) and -$\log_{10}$ adjusted p-value (y-axis). Distinct genes are highlighted in red. (**E**) RNA-seq TPM values of selected HALLMARK EMT pathway genes from EtOH- and Dex-treated p63KO cells, and siNON- and siGATA2-treated p63KO cells. Bar graphs represent mean ±SD, n=3. (**F**) (upper) Relative cell proliferation in siNON- and siGATA2-treated p63KO cells in the presence or absence of Dex treatment. Data is normalized to day 0 and represents the mean ±SD of 10 images per day. (lower) Summary table of the statistical comparison of between siNON and siGATA2 exposed p63KO cell proliferation at the three last timepoints. (**G**) 3D gel invasion of CTRL and p63KO cells after day 6 of EtOH, Dex, or K-7174 treatments. Two of the highest invasion points were utilized (n=16-30). Statistical significance in cell proliferation line graph was calculated with Two-way ANOVA with Bonferroni post hoc test. Statistical significance in box plot was calculated with One-way ANOVA with Bonferroni post hoc test. *, $p< 0.05$; **, $p< 0.01$; ***, $p< 0.001$. PRAD, prostate adenocarcinoma; TCGA, the cancer genome atlas.

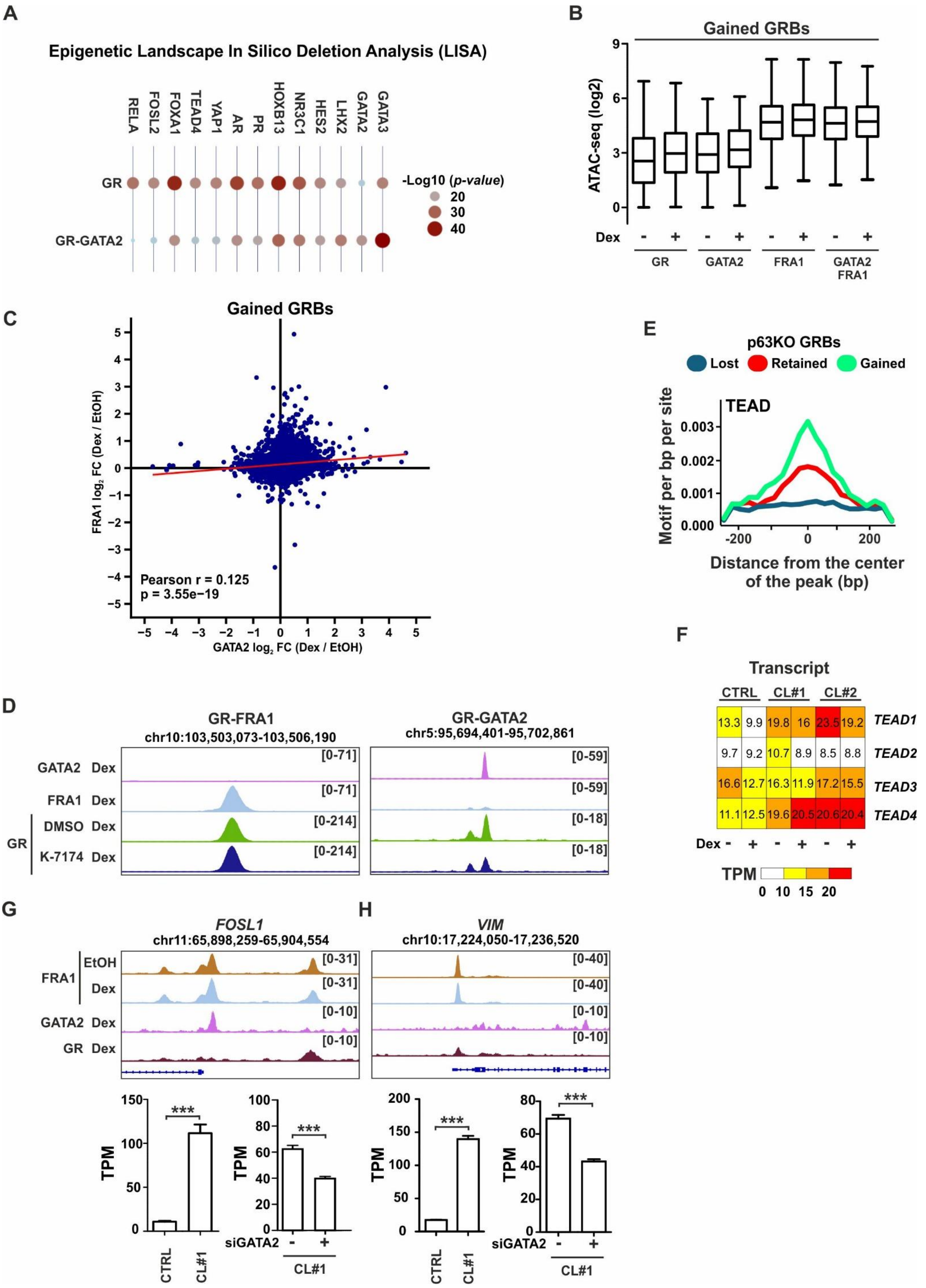

A
Epigenetic Landscape In Silico Deletion Analysis (LISA)
RELA
FOSL2
FOXA1
TEAD4
YAP1
AR
PR
HOXB13
NR3C1
HES2
LHX2
GATA2
GATA3
GR
GR-GATA2
-Log10 (p-value)
20
30
40
B
Gained GRBs
ATAC-seq (log2)
9
6
3
0
Dex
- + - + - + - +
GR
GATA2
FRA1
GATA2 FRA1
C
Gained GRBs
FRA1 log2 FC (Dex / EtOH)
GATA2 log2 FC (Dex / EtOH)
Pearson r = 0.125
p = 3.55e−19
E
p63KO GRBs
Lost
Retained
Gained
TEAD
Motif per bp per site
0.003
0.002
0.001
0.000
-200
0
200
Distance from the center of the peak (bp)
F
Transcript
CTRL
CL#1
CL#2
13.3 9.9 19.8 16 23.5 19.2 TEAD1
9.7 9.2 10.7 8.9 8.5 8.8 TEAD2
16.6 12.7 16.3 11.9 17.2 15.5 TEAD3
11.1 12.5 19.6 20.5 20.6 20.4 TEAD4
Dex - + - + - +
TPM
0 10 15 20
D
GR-FRA1
chr10:103,503,073-103,506,190
GR-GATA2
chr5:95,694,401-95,702,861
GATA2 Dex
FRA1 Dex
GR
DMSO Dex
K-7174 Dex
[0-71]
[0-71]
[0-214]
[0-214]
[0-59]
[0-59]
[0-18]
[0-18]
G
FOSL1
chr11:65,898,259-65,904,554
FRA1
EtOH
Dex
GATA2 Dex
GR Dex
[0-31]
[0-31]
[0-10]
[0-10]
H
VIM
chr10:17,224,050-17,236,520
[0-40]
[0-40]
[0-10]
[0-10]
TPM
***
CTRL
CL#1
siGATA2
- +
CL#1

**Supplementary Figure S12. GATA2 and FRA1 modulate GR action and transcriptional changes.** (**A**) Prediction of TF activity at p63KO GRBs with or without GATA2 binding using epigenetic Landscape In Silico Detection Analysis (LISA). Data depicted as $-\log_{10}$ p-value. (**B**) $\log_2$ tag enrichment of ATAC-seq in p63KO cells at p63KO-gained GRBs, categorized by overlap with only GATA2, only FRA1, both (GATA2 FRA1) or neither (GR). (**C**) Scatter plot show Dex-mediated $\log_2$ fold change ($\log_2$ FC) of EtOH and Dex treatments (Dex / EtOH) of (x-axis) GATA2 and (y-axis) FRA1 binding at gained GRBs. Pearson correlation coefficient and p-value are shown in the plot. (**D**) Genome browser track of GATA2, FRA1, and K-7174 or DMSO treated GR ChIP-seq from p63KO cells at p63KO-gained GRB loci containing either GR-FRA1 (left) or GR-GATA2 (right) peaks. (**E**) Motif density of TEAD at indicated p63KO GRBs. Histograms represent ± 240 bp around the center of the peak, and enrichment intensity represents motif per bp per site. (**F**) Heatmap of TEAD-family member TPM values in CTRL and p63KO cells. (**G-H**) Genome browser track (upper) of GATA2, GR, and FRA1 ChIP-seq from p63KO cells at (**G**) *FOSL1* or (**H**) *VIM* locus. Bar plot (bottom) of corresponding RNA-seq TPM values of (**G**) *FOSL1* or (**H**) *VIM* expression in CTRL and p63KO cells and in siNON- and siGATA2-treated p63KO cells. Bar graphs represent mean ±SD, n=2-3.
